# Multi-barrier unfolding of the double-knotted protein, TrmD–Tm1570, revealed by single-molecule force spectroscopy and molecular dynamics

**DOI:** 10.1101/2025.08.27.672562

**Authors:** Fernando Bruno da Silva, Szymon Niewieczerzal, Iwona Lewandowska, Mateusz Fortunka, Maciej Sikora, Laura-Marie Silbermann, Katarzyna Tych, Joanna I. Sulkowska

## Abstract

The doubly knotted motif is one of the least expected features in proteins, occurring in both globular and transmembrane forms. Here, we focus on globular protein members of the methyltransferase family: the TrmD–Tm1570 protein, which contains two deep 3_1_ knots, and the single-knotted proteins TrmD and Tm1570, all from *Calditerrivibrio nitroreducens*. Using various biophysical experimental techniques and computer simulations with AI-based methods, we studied their thermal and thermodynamic stability, as well as their mechanical unfolding. Based on molecular dynamics (MD) simulations, with the Structure-Based C*α* Model (SBM-C*α*) and UNRES (coarse-grained), we show that native contacts alone are not sufficient to fold double-knotted proteins. However, native contacts are sufficient to fold the single-knotted proteins TrmD and Tm1570 into their native conformations. Using the same model, we identified four possible unfolding and untying pathways, in which each domain can self-tie independently at some stage of the process. Optical tweezers (OT) experiments show that this process is also reversible, although the stretched state remains knotted. In addition, we observed higher thermal and mechanical stability in Tm1570 compared with TrmD, which is partly attributable to the position of the knot core. Overall, our results suggest that double-knotted protein from the SPOUT family can only partially self-fold, and that full knotting may require the assistance of a chaperone.

## Introduction

Understanding the relationship between a protein’s sequence and its structure remains one of the most important challenges in structural biology (***Dobson, 2003***). Despite differences in their sequences, proteins preserve certain structural features known as motifs. Among these, entanglement is a comparatively recent discovery and is starting to be recognized as important, due to potential benefits, such as enzymatic activities (***Nureki et al., 2004***; ***Perlinska et al., 2020a***), ligand binding (***Wagner et al., 2007***; ***Christian et al., 2016***; ***Perlinska et al., 2020b***), contribution to reactivity or other electronic structure properties (***Alves Silva et al., 2024***), introduces thermal ((***Silva et al., 2022***; ***Sayre et al., 2011***)) or mechanical stability ((***Sułkowska et al., 2008***; ***Liu et al., 2017***; ***Dzubiella, 2009***)), that a non-trivial topology can bring (***Schaufelberger, 2020***; ***Ashbridge et al., 2022***; ***Scholl and Deniz, 2022***). One type of entanglement found in proteins is a knot formed along the backbone. There are a few types of knots verified with crystal structures, based on the KnotProt Database: 3_1_, 4_1_, 5_2_, 6_1_ (***Mansfield, 1994***; ***Bölinger et al., 2010***), the recently discovered 7_1_ knot (***Hsu et al., 2024***) and the double-knotted 3_1_#3_1_ (***da Silva et al., 2023***; ***Perlinska et al., 2024***). There might be more types and many knotted families, as shown by structures predicted by AlphaFold. Right now, the AlphaKnot 2 database has over 600,000 knotted structures (***Niemyska et al., 2022***).

Here, we focus on a double-knotted motif (***Perlinska et al., 2024***; ***Brems et al., 2022***), a knot composed of two individual trefoil knots, since their impact on the energy landscape is very interesting and has not yet been studied. This structural motif is found in several families of proteins whose proper function is crucial to the cell. In addition, thanks to advances in structural biology and machine learning, this motif can also be artificially constructed to form helical structures (***Doyle et al., 2023***), and formed from the combination of individually knotted proteins (***Sayre et al., 2011***). However, the doubly knotted knot is unique in terms of its folding or unfolding mechanism. All known single knots found in proteins are categorized as twisted knots, which means that they need just one threading event during folding in order to reach the native state. In contrast, the double knotted protein requires two threading events and each of these threading events constitutes a topological barrier.

As stated above, there is only one solved structure of a naturally occurring double knotted protein (PDB ID 881N); however, there are several protein families which are predicted to embed a composite 3_1_#3_1_ knot (***Perlinska et al., 2024***). In total, there are more than 1200 predicted proteins with double knot, including globular and transmembrane proteins. All of the double knotted structures found by (***Perlinska et al., 2024***) display a composite 3_1_#3_1_ knots. Since the trefoil knot is the most common knot type found in proteins (***Dabrowski-Tumanski et al., 2019***), we expected to see composite knots containing the 3_1_ knot (***Niemyska et al., 2022***). Analysis of a large dataset (nearly 1,000 proteins) identified five domain arrangements with a doubly knotted structure. The double knot topology is found in membrane proteins from the CaCA family (UniProtKB ID: A0A179I9N9), which function as ion transporters; Carbonic anhydrase (UniProtKB ID: A0A0B7AKD5), which catalyzes the hydration of carbon dioxide; and from three domains from the SPOUT superfamily (UniProtKB ID: Q4DMW6, A0A498KD62 - Nep1-Nep1, and E4THH1 - TrmD-Tm1570), the largest group of deeply 3_1_ knotted proteins (***Sulkowska, 2020***).

The protein with a confirmed double-knotted topology, TrmD-Tm1570 belongs to the SPOUT family (***Figure 1***A). It has previously been shown that the TrmD domain from TrmD-Tm1570 from *Calditerrivibrio nitroreducens* (*Cn*) is biologically active and has the same activity as TrmD from a different organism, i.e., m1G37-tRNA methylation (***Perlinska et al., 2020a***). However, the function of the Tm1570 domain from TrmD-Tm1570 remains unclear, requiring further investigation into both the individual protein and its TrmD-Tm1570 activity. The degradation process studies of single-knotted proteins have shown that entangled proteins can be degraded or can impairs protein degradation by ATP-dependent proteases (***San Martín et al., 2017***; ***Sriramoju et al., 2018***; ***Sivertsson et al., 2019***; ***Fonseka et al., 2021***; ***da Silva et al., 2023***), thus, this is still an area of investigation. It was also showed that the barrel-shaped ATP-dependent ClpXP protease can degrade TrmD-Tm1570, TrmD, and Tm1570 (***da Silva et al., 2023***). However, SDS-PAGE analysis revealed the presence of a fraction of TrmD-Tm1570, indicating incomplete degradation by ClpXP.

**Figure 1.**
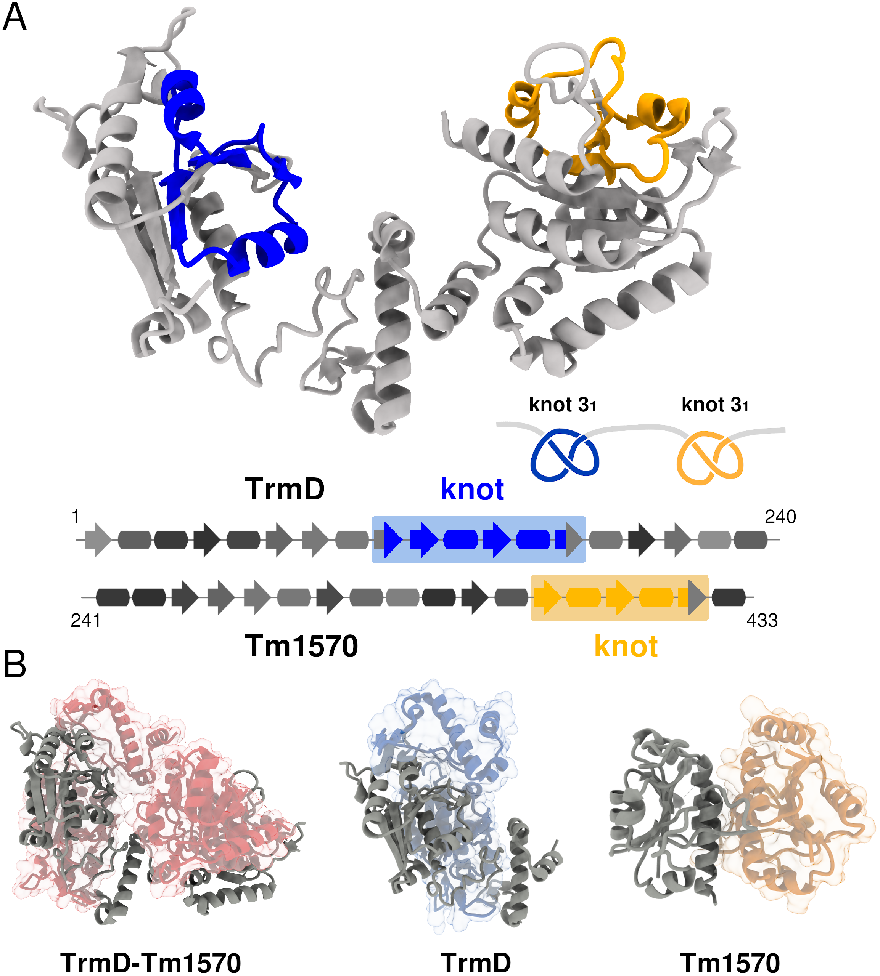
(A) Crystal structure of TrmD-Tm1570 monomeric protein (Protein Data Bank ID code: 8B1N). The double trefoil-knot, 3_1_#3_1_, is shown in blue for the TrmD domain and in yellow for the Tm1570 domain. The secondary structure representations for TrmD, residues 1-240, and Tm1570, residues 241-433, are represented in the bottom of panel A. The *β*-sheets and *α*-helices are represented by rectangles and arrows, respectively. (B) Dimeric crystal structures of TrmD-Tm1570 (PDB ID: 8B1N), TrmD (PDB ID: 8BYH), and Tm1570 (PDB ID: 8RI0).

Herein, our goal is to elucidate the (un)folding processes of double knotted protein and stabilising interactions by different approaches. Understanding the folding pathways and cellular degradation of double-knotted proteins is essential for uncovering their biological roles and potential applications. Because unfolding, whether induced by heat or mechanical stress, lies at the heart of both processes, we concentrate here on thermal and mechanical unfolding, combining experimental and theoretical approaches. In addition, the double-knotted enzyme TrmD-Tm1570 has proven to be extremely labile: every thermal unfolding/refolding attempt leads to aggregation. We therefore place particular emphasis on mechanical unfolding, a technique so far applied only to singly knotted systems (***Arai et al., 1999***; ***Ziegler et al., 2016***; ***Rivera et al., 2023***).

We focus on the deeply double knotted, TrmD-Tm1570, and single knotted TrmD and Tm1570, proteins from *Calditerrivibrio nitroreducens* (*Cn*). Molecular dynamics simulations and experimental methods were conducted to investigate the domains and knot stability, and potential (un)folding pathways. First, the folding pathway for three proteins was investigated with various coarse-grained models. Next, the unfolding process by thermal unfolding simulations was carried out using the Structure-Based C*α* Model (SBM-C*α*) in order to describe the (un)folding process of all three proteins. Second, protein pulling was conducted experimentally by optical tweezers, and by a mechanical simulation under constant velocity for the purpose of investigation. All atomic simulations in an explicit solvent were additionally performed to determine the position of the protein terminus, which facilitates the interpretation of experimental data. The experimental data was also processed using AI methods. Thermal resistance analysis was estimated based on differential scanning fluorimetry (DSF). This combined approach offers valuable insights into the mechanical properties, stability, and unfolding mechanism of the first characterized double knotted protein, TrmD-Tm1570.

## Results

### Double knotted fusion protein, TrmD-Tm1570

TrmD-Tm1570 (PDB ID: 8B1N) is a protein consisting of two subunits: TrmD (residues 1-240) and Tm1570 (residues 241-433), ***Figure 1***A. The single domains, Tm1570 and TrmD, are homodimeric proteins and their active sites are located in the knot region; both structures contain a deep 3_1_-trefoil knot (***Figure 1***B). The TrmD domain contains seven *β* strands and nine *α* helices. The Tm1570 subunit has five *β* strands and eight *α* helices. In both domains, there are five *β* strands sandwiched between *α* helices on either side forming a well-defined structural motif. TrmD is the larger protein with 240 residues and the knot core is located between residues 85 - 129. On the other hand, Tm1570 is shorter, with 193 residues, and the knot core ranges from residue 113 to 158, ***Figure 1***B. Binding sites and general details of TrmD-Tm1570 are described in ***da Silva et al. (2023)***.

### Double-knot protein cannot be knotted in a known structure-based C*α* model

One intriguing question about double-knotted proteins is whether they can self-tie. The TrmD-Tm1570 protein belongs to SPOUT family. It has previously been shown by many groups with different types of structure-based models that members of this family which possess a single knot, such as Yibk and Yeba, can self-tie (***Sułkowska et al., 2009)***. However, adding non-native contacts makes this process easier (***Wallin et al., 2007)*** also in the case of other knotted proteins (***Škrbić et al., 2012)***. It was suggested that the knot in Yibk is formed via the so-called twist pathway (***Wallin et al., 2007***; ***Sułkowska et al., 2009)***, where the shorter terminal is threaded across a twist loop (almost in the native position). This implied that, in the case of a double knot, the knot can be formed either from the C-terminus (threading 40 residues) or from the N-terminus (threading 50 residues).

Based on the structure-based C*α* model (SBM-C*α*) we found that both TrmD and Tm1570 can self-tie via threading of the C-terminus, however the success rate is very low (as was the case for other SPOUT members) (***Sułkowska et al., 2009)***. We used the same model (and several different versions of it with different maps of native contacts (cut-off and high frustrate contacts) and shapes of attractive potential) for double-knotted TrmD-Tm1570 protein and did not find a single trajectory in which two knots were formed in the native position. In this case, the number of simulations was twice (and much longer) that performed for single proteins, TrmD and Tm1570, and we tested different temperatures. We also used a different SBM-C*α* models developed by the Cieplak group, and a completely different type of model UNRES (***Czaplewski et al., 2018)***. None of the models allowed the protein to fold into its native state with a double knot. One possible explanation is that the protein needs support from the ribosome to tie a knot, as has previously been shown for a protein with a very deep knot (***Dabrowski-Tumanski et al., 2018)***. Additionally we can speculate that double-knotted proteins may need cellular factors such as chaperones (***Mallam and Jackson, 2012)*** to refold to the correct knotted native state.

Thus, to understand how a protein with 3_1_#3_1_ topology can double knot, we investigated the unfolding and unknotting pathways with the same SBM-C*α* model (see Methods and Materials - Mechanical unfolding simulations).

### Double-knot can be unfolded and unknotted in a SBM-C*α*

Analysis of the simulated unfolding trajectories of TrmD-Tm1570 shows that the unfolding process of this protein can be realized in several ways, ***Figure 2***. Based on the simulations, we observed four predominant unfolding and untying pathways, ***Figure 3***. These set of simulations were obtained as follows: (i) trajectories where TrmD unfolds first and its knot unties before Tm1570 unfolds; (ii) trajectories where Tm1570 unfolds and unties first, followed by TrmD; and (iii) trajectories where TrmD unfolds first, then Tm1570, after which the TrmD knot unties and finally the Tm1570 knot unties.

**Figure 2.**
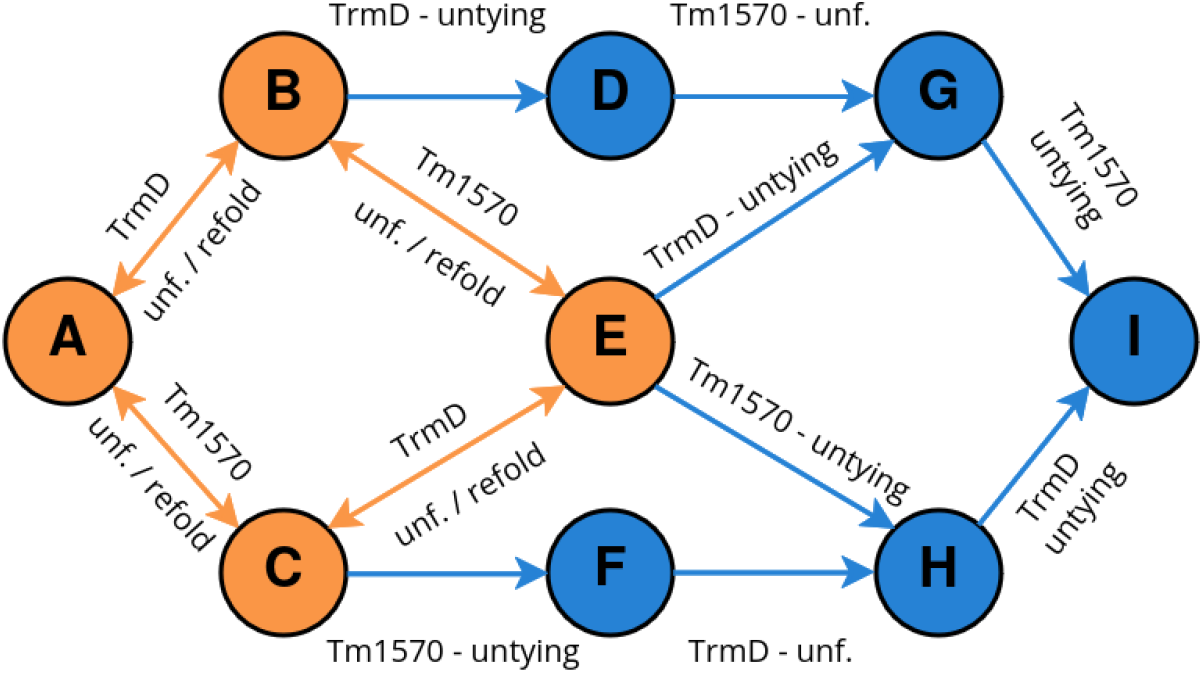
Schematic representation of the unfolding mechanism of the double-knotted protein TrmD-Tm1570. The figure shows possible unfolding pathways observed in the structure-based model simulations. Starting from a native double-knotted topology (A), the system undergoes a series of intermediate processes (B, C, D, E, F, and G) before reaching its unfolded state (I), which is represented by a random unknotted/trivial conformation. Orange and blue colors represent the reversible and non-reversible processes, respectively.

**Figure 3.**
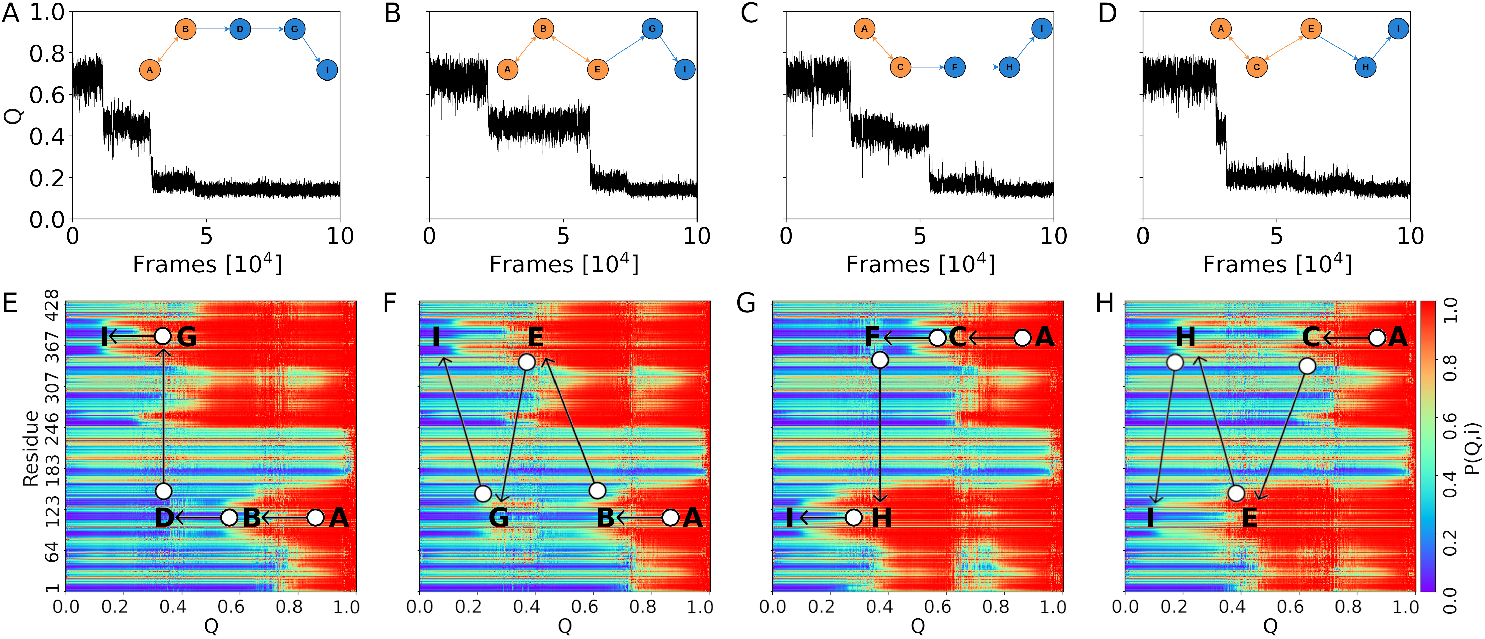
The unfolding trajectories and the probabilities of contact formation per residue. A, B, C, D display the fraction of native contacts, *Q*, as function of the number of frames. E, F, G, and H show the probability of residue *i* being in contact with its native contacts as function of *Q*. Panels A/E, B/F, C/G, and D/H correspond to the unfolding pathways 1, 2, 3, and 4, respectively. **Figure 3—figure supplement 1**. Unfolding trajectories observed less frequently, rare events. **Figure 3—figure supplement 2**. Refolding dynamics of a fusion protein, visualized through snapshots at key time steps **Figure 3—figure supplement 3**. Unfolding trajectory and knot dynamics of the TrmD–Tm1570 from panel A.

The schematic representation in ***Figure 2*** of double knotted native (N) state is shown as 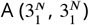, there is a series of intermediate states in 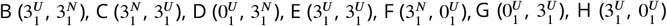, and an unfolded (U) and unknotted state in 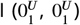. The indices *N* and *U* are related to the native and unfolded state, respectively. In parentheses, we provide information about the topology of TrmD and Tm1570 domains, respectively, and the superscript indicates whether a domain is in the native (N) or unfolded (U) state. The pathways, as described before, in the course of simulations are: pathway 1 - A↔B → D → G → I; pathway 2 - A↔B↔E → G → I; pathway 3 - A↔C → F → H → I; and pathway 4 - A↔C↔E → H → I, ***Figure 3***A, B, C, and D, respectively. In the case of path 1 and 2, the knot unties in the N-terminal domain (TrmD) first, and then in the C-terminal domain (Tm1570). On the other hand, in the case of path 3 and 4, first the knot unties in the C-terminal domain (Tm1570), and then in the N-terminal domain (TrmD).

Pathway 1 (A↔B → D → G → I) starts with a reversible step (unfolding/refolding, A↔B) of the TrmD domain. Once the TrmD domain unfolds, the knot untying event is observed, B→D. Next, by a non-reversible process, the Tm1570 domain unfolds and unties (D→G→I). Here we observe two interesting cases in which the knot topology influences protein unfolding - according to how deep a knot is along the chain. In the case of the TrmD domain, we are likely to observe refolding of the domain since the untying of the knot is restricted. This knot possesses a very long C-terminal tail including the Tm1570 domain. The only way to untie the TrmD knot is from the N-terminal tail, however, this tail consists of 50 residues. Thus, the secondary structure in the N-terminal region must unfold first. As a consequence of the knot remaining in its native position, the TrmD domain can fold and unfold around the knotted core within thermal fluctuations. At some point, when the N-terminal tail is completely unfolded, the knot can spontaneously untie and unfold itself (***Figure 3***E).

The knot in the Tm1570 domain is shallower than that in TrmD, and less constrained by the secondary structure.

In pathway 2, (A↔B↔E → G → I) is described by two reversible processes, A↔B and B↔E, in which the TrmD and Tm1570 domains can both unfold/refold. However, the refolding process is observed only when the knot location along the chain is close to its native position (***Figure 3 - figure Supplement 1*** and ***figure Supplement 2***). Here, the TrmD domain unties first (E→G), followed by the untying of the Tm1570 domain (G→I), where I is the unfolded state.

In the case of pathway 3 (A↔C → F → H → I) and pathway 4 (A↔C↔E → H → I), ***Figure 3***C and D, the process starts with the unfolding of the Tm1570 domain (A↔C). This pathway is followed by Tm1570 untying (C→F). From state F to unfolded state, I, the TrmD domain unfolds and is untied through an irreversible process. In pathway 4, the unfolding of Tm1570 is followed by the reversible unfolding of the TrmD domain (C↔E). The unfolded state, I, is reached by two consecutive untying processes: first of the Tm1570 knot (E→H) and then the TrmD knot (H→I).

### Detailed Insight into untying process

Examples of TrmD-Tm1570 unfolding trajectories representing each of the four main unfolding pathways are presented in ***Figure 3***A-D showing the time dependence of the fraction of remaining native contacts (Q). Based on the obtained unfolding trajectories, we calculated the probability of contact formation per residue, *P* (*Q, i*), for each pathway, see ***Figure 3***E-H. The *P* (*Q, i*) shows which region of the protein unfolds as a function of *Q*.

In pathways 1 and 3, the unfolding process starts with the unfolding of an individual domain directly followed by its untying. This is followed by the unfolding and untying of the other domain, after which the protein reaches the fully unfolded state. In pathway 1, TrmD unfolds and unknots first, while Tm1570 is the first to unfold and unknot in pathway 3, see ***Figure 3***A/E and ***Figure 3***C/G, respectively.

In pathways 2 and 4, we observe both domains unfolding first, and subsequently unknotting. TrmD unfolds first in pathway 2, and Tm1570 in pathway 4, see ***Figure 3***B/F and ***Figure 3***D/H, respectively. For these pathways, we observe the knot translocation in the already unfolded domain during the unfolding of the other domain. In the case of pathway 2, the knot located in the TrmD domain slides towards the N-terminus during the Tm1570 unfolding process, B↔E→G (0.4 < *Q* < 0.6), see ***Figure 3***F. In the case of pathway 4, the knot located in the Tm1570 domain slides in the direction of the C-terminus during the TrmD unfolding process, C↔E→H (0.4 < *Q* < 0.6), see ***Figure 3***H.

Pathway 2, with TrmD unfolding before Tm1570, is the most populated one. This finding is in agreement with previous results on the degradation process of the TrmD-Tm1570, which have shown that Tm1570 has higher stability than TrmD. The same result was observed, experimentally and computationally, for the degradation rate of the individual domains in (***da Silva et al., 2023)***. Knot depth, in both the fusion protein and in the individual domains, is the key determinant of stability of the proteins, since the knot core from Tm1570 is located closer to the C-terminus than the knot core from TrmD.

### Mechanical unfolding in optical tweezers experiments

Constant velocity force-extension measurements were collected for Tm1570, TrmD and the Tm1570-TrmD fusion protein (***Figure 4)***. Measurements of unfolding and refolding traces showed a consistent unfolding pathway for TrmD, with one smaller unfolding event with an increase in contour length of 10.6 nm (+/- 1 nm) and one larger unfolding event with an increase in contour length of 35.3 nm (+/- 1.5 nm) giving a total change in contour length of 45.8 nm (+/- 1.8 nm). All the values between parentheses correspond to the standard deviations of the lengths.

**Figure 4.**
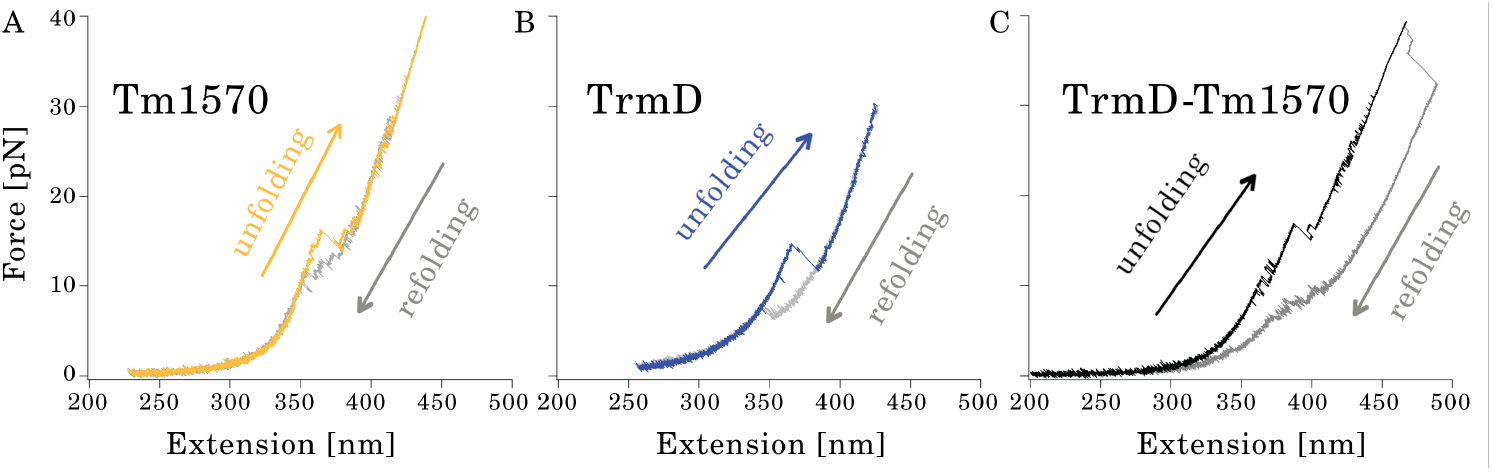
Experimental unfolding and refolding of Tm1570, TrmD and the Tm1570-TrmD Fusion using single-molecule optical tweezers. In each trace the unfolding is colored (orange for Tm1570, dark blue for TrmD and black for the Fusion), while the refolding trace is in grey. The traces were measured at a constant velocity of 20 *nms*^−1^ for both the retraction and approach segments (unfolding and refolding). **Figure 4—figure supplement 1**. Experimental unfolding under constant velocity of 20 and 500 *nms*^−1^.

For Tm1570, a variety of different unfolding pathways were observed. Overall, however, the most commonly observed pathway exhibited three main unfolding events, where the changes in contour length added up to a total of 53.3 nmon average (+/- 3.8 nm). The expected changes in contour length for each domain are given by Δ*L*_*C*_ = (*n*_*aa*_ ∗0.365 nm)-*l*_*init*_, where *n*_*aa*_ is the number of amino acids between the two points where the force is applied, 0.365 nm is the average length of an amino acid and *l*_*init*_ is the initial distance between the two points where the force is applied, in nm. For example, for a protein that is unfolded from the N- and C-termini, has an initial N- to C-terminus distance of 2.35 nm and has 191 residues between the attachment points as is the case for Tm1570, the expected change in contour length from the fully folded state to the fully unfolded state is Δ*L*_*C*_ =(191*0.365 nm)-2.35 = 67.4 nm. The measured changes in contour length for all three proteins suggest the presence of a structure or structures which are not unfolded under the applied forces at these constant stretching velocities, which we attribute to the presence of the knots in the structure. A summary of the percentage of traces showing complete unfolding and the corresponding contour-length changes for all three proteins is provided in ***Table 1*** and ***Table 2***.

**Table 1.** Percentage of traces showing (near) complete unfolding for TrmD, Tm1570, and TrmD-Tm1570.

| Protein | Percentage (%) |
| --- | --- |
| TrmD | 100 |
| Tm1570 | 75 |
| TrmD-Tm1570 | 45 |

**Table 2.** Contour length changes for fully unfolded traces for TrmD, Tm1570, and TrmD-Tm1570. ^1^No. - Number of unfolding traces.

| Protein | Avg (nm) | Std (nm) | No.<sup>1</sup> |
| --- | --- | --- | --- |
| TrmD | 45.8 | 1.8 | 38 |
| Tm1570 | 53.9 | 4.0 | 16 |
| TrmD-Tm1570 | 89.3 | 4.5 | 22 |

Our measured changes in contour length are in good agreement with the simulations (***Figure 5)***. For Tm1570, the change in length obtained in the simulation is (54.37 nm - 2.35 nm) = 52.02 nm, c.f. the 53.3 nm we measure experimentally on average, with the most common pathway in simulations showing steps of 2.38 nm, 7.30 nm, 28.01 nm and 14.13 nm while experimentally we typically observed steps of 13 - 15 nm, 25 - 30 nm and 9 - 13 nm (likely representing a sum of the first two events observed in the simulation). For TrmD, the observed change in length in the simulations is (62.14 nm - 4.68 nm) = 57.46 nm, while experimentally we observe an average length change of 45.8 nm. However, the first unfolding steps in the simulation involve a likely very mechanically weak three-helix bundle at the C-terminus of the protein (see steps 1 and 2 in ***Figure 5***D). Assuming that this structure is either unfolded at a force below the sensitivity of the optical tweezers, or is not folded in the monomeric state of the protein, the remaining unfolding steps correspond very well to the experimental data, with the unfolding steps seen in the simulations of 9.08 nm, 30.31 nm and 2.94 nm adding up to 42.33 nm, and our measured change in contour length being 45.8 nm (+/- 1.8 nm) where, most likely, the final two steps in the simulation combine into a single event in the experimental traces.

**Figure 5.**
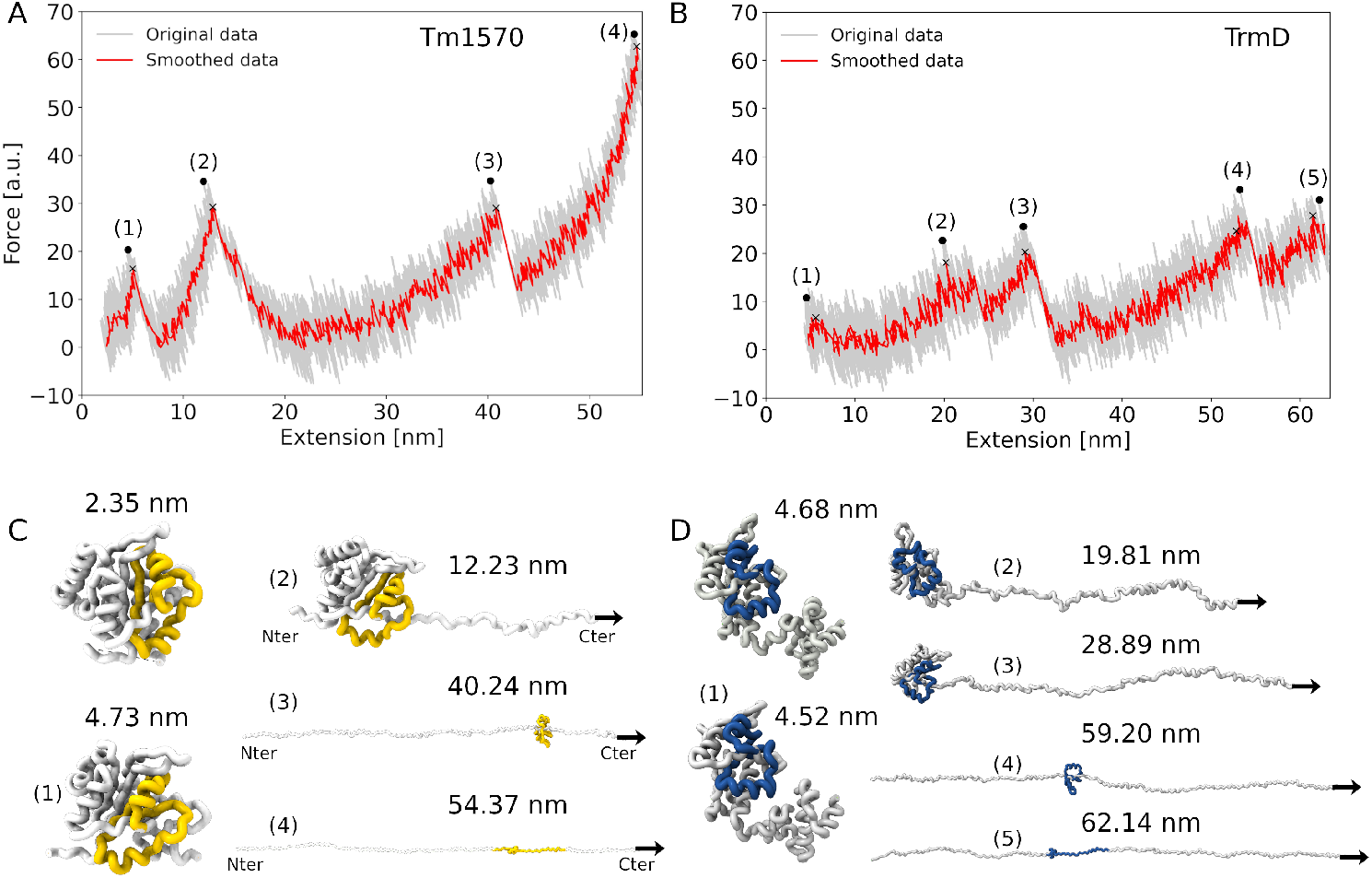
Force versus extension of Tm1570 (A) and TrmD (B). The original data collected from the simulation are shown in light grey and the smoothed data by moving average algorithm in red. Each peak is represented by an index. Panels C and D show cartoon representations of Tm1570 and TrmD native structures and snapshots from the simulation. The indices (1, 2, 3, and 4) and (1, 2, 3, 4, and 5) correspond to the same indices in A and B, respectively. The knot core is highlighted in yellow and in blue for the Tm1570 and TrmD domains, respectively. **Figure 5—figure supplement 1**. Clustering of the unfolding trajectories for Tm1570 obtained via SOM analysis. **Figure 5—figure supplement 2**. Clustering of unfolding trajectories for TrmD obtained via SOM analysis. **Figure 5—figure supplement 3**. Distribution of simulations for each protein across clusters **Figure 5—figure supplement 4**. End-to-end cysteine distances, all atoms MD - probability density and free energy landscapes for TrmD, Tm1570, and the TrmD–Tm1570.

### Stability of knot motifs at low pulling speeds in simulations

In order to get a deeper understanding of the mechanical unfolding mechanism, especially to unveil the way in which knots tighten during this process, we performed MD simulations mimicking the optical tweezers-based pulling experiments. A computational approach is necessary because the experimental methods do not give insight at a comparable level of resolution. We considered three different speeds at which the protein was unfolded: 0.05 Å/*τ*, 0.10 Å/*τ*, and 0.15 Å/*τ*.

Force-extension curves obtained at a pulling velocity of *ν* equal to 0.05 Å/*τ* for TrmD, Tm1570, and TrmD-Tm1570 proteins, respectively, are shown in ***Figure 5***A-B and ***Figure 6***A. In general, the stretching of each protein produces various force-extension profiles. However, it is possible to select the most typical, commonly occurring patterns. The presented curves are representative force-extension traces located in the most populated cluster. These selections are based on the SOM cluster analyses (SOM is described in detail in the *Methods Section*). In this method, after identifying the main cluster, we obtained the corresponding cluster centroid and then determined the nearest simulation point (***Figure 5*** and ***Figure 6)***.

**Figure 6.**
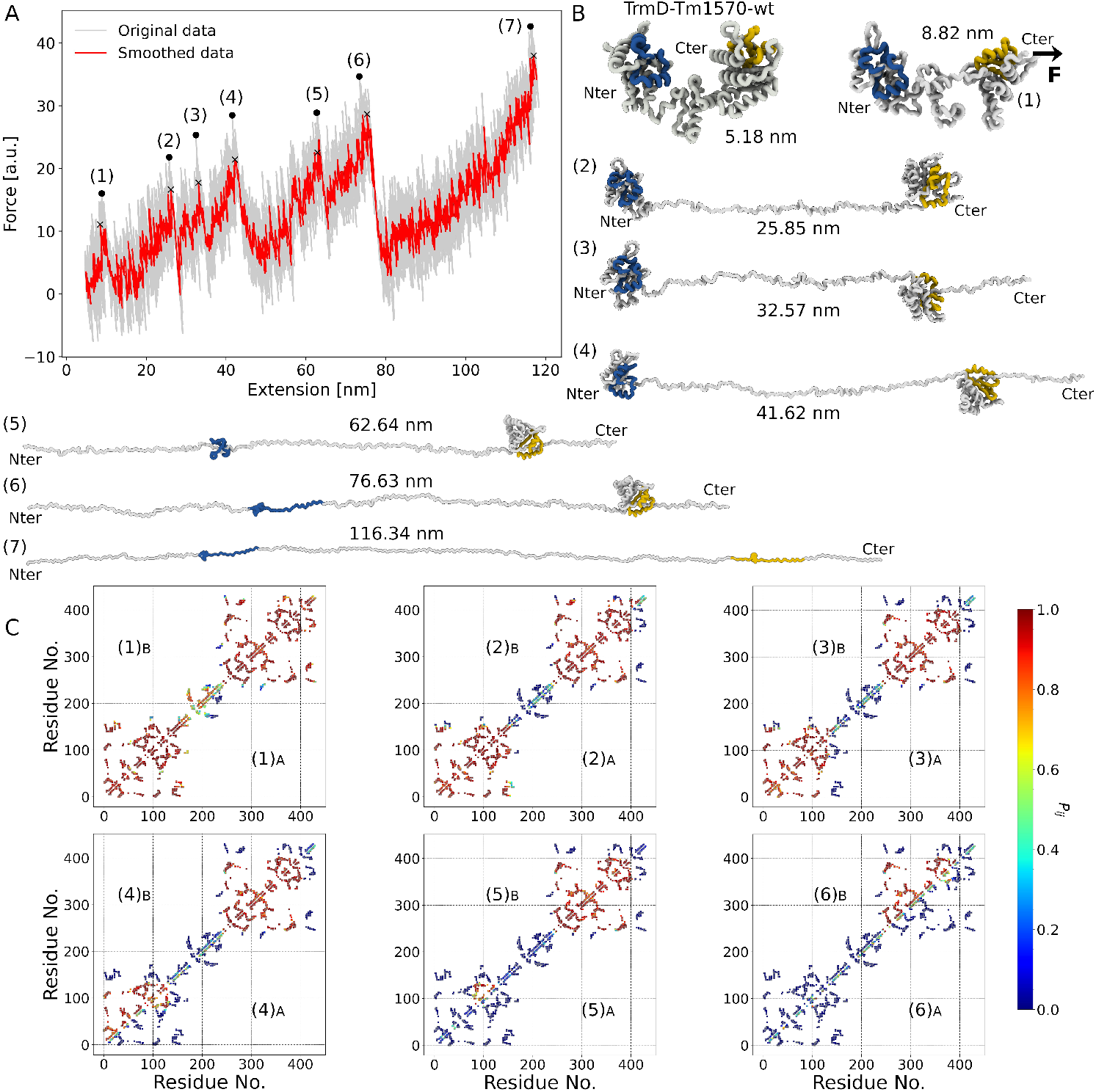
(A) Force versus extension of TrmD-Tm1570 protein. The original data from the simulation is shown in light grey and the smoothed data by moving average algorithm in red. Each peak is represented by an index. (B) Cartoon representation of the TrmD-Tm1570 fusion protein’s native structure and snapshots from the simulation. The indices 1, 2, 3, 4, 5, 6, and 7 correspond to those in panel A. The knot core is highlighted in blue and yellow for TrmD and Tm1570, respectively. The knot core length for all conformations is always pictured as encompassing the same number of residues as that in the native structure for clarity. (C) Probability of native contacts - upper and lower triangle based on 1000 conformations before (B) and after (A) of each index, respectively, where red points indicate a high probability of there being a contact and blue points indicate a low probability of there being a contact (see colour map). **Figure 6—figure supplement 1**. Clustering of unfolding trajectories for Tm1570-TrmD obtained via SOM analysis.

In the case of Tm1570 mechanical unfolding, there are four distinct peaks, or rupture points, see ***Figure 5***A. All of the force-extension curves and cluster distributions can be found in Figure S2 and S5A in Supporting Information. Snapshots of representative conformations at each peak are presented in ***Figure 5***C. The end-to-end Tm1570 native distance is equal to 2.35 nm, however, the all-atom an explicit solvent MD simulations in equilibrium showed that this value can fluctuate up to 3.0 nm. Peaks 1, 2, and 3 correspond to intermediate states (***Figure 5***C(1-3)). The last peak, number 4, corresponds to the maximal extension, according to the criteria adopted during the analyses (*Q*_c_ = 5% *Q*_native_). The end-to-end distance for the maximum extension is equal to 54.37 nm, which is in good agreement with the experimental results.

To understand what the unfolding pathway looks like, when the knot tightens, and how a knot influences the process, we analysed the location of the force peaks and the corresponding conformations. The end-to-end distances to the peaks 1, 2, and 3 are equal to 4.73 nm, 12.23 nm, and 40.24 nm, respectively. Similarly to thermal unfolding, Tm1570 first unfolds from the C-terminus (rupture points 1 and 2), without deformation of the knot. Next, the N-terminal end is extended (rupture point 3) which finally causes a tightening of the knot (rupture point 4), see ***Figure 5***. During the mechanical unfolding process of Tm1570, the knot core displays a high stability compared with other regions of the protein.

***Figure 5***B shows a representative force-extension curve for the mechanical unfolding of TrmD, selected based on SOM analysis (for details see Figure S3 and S5B in the Supporting Information). The native end-to-end distance is equal to 4.68 nm, in contrast, equilibrium all-atom an explicit solvent MD simulations revealed that this value can vary by as much as 5.0 nm, and there are five representative peaks which correspond to the intermediate processes to reach the complete elongation of the protein. Simulations show that TrmD has a highly flexible region located in its C-terminus, which is responsible for binding tRNA. In organisms in which TrmD is not fused with Tm1570, the C-terminus is stabilised by the interface formed by dimerization. The first peak does not correspond to the elongation of the chain, but is a result of the force applied to disrupt the native conformation of the protein, ***Figure 5***D(1) and S7 in the Supporting Information. The end-to-end distance of the first point of rupture is equal to 4.52 nm. Next, one can observe that the elongation of the C-terminus region until the chain up to the portion that is involved in the knot core formation is unfolded (rupture point 2, 19.81 nm). Subsequently, the protein starts to unfold from the N-terminus, where the chain loses almost all tertiary and secondary structure up to the knot core (rupture point 3). From that moment both total unfolding of the N-terminal region and knot tightening take place (rupture point 4 up to 5). In the same manner as Tm1570, the last two points of rupture are related to the processes of tightening the knot in both proteins. Therefore, the Tm1570 and TrmD unfolding mechanisms are similar, and the process can be described by three steps as follows: first, C-terminal region elongation; second, N-terminal region elongation; and third, knot tightening and full elongation. However, despite their similarities, the force required to rupture each event is different. Following the criteria, *Q*_c_, the force required, *F*, to observe the complete elongation and knot tightening is twice as high for Tm1570 as for TrmD. Thermal and mechanical unfolding follow different reaction coordinates, but nevertheless both approaches show Tm1570 to be more stable than TrmD.

Next, we show how the fusion of TrmD and Tm1570 changes the unfolding pathways of both domains in TrmD-Tm1570. Herein, again based on the SOM approach, we clustered unfolding pathways and used the same criteria to analyse curves. A representative force-extension curve for TrmD-Tm1570, mechanical unfolding, is presented in ***Figure 6***A and, as expected, it is much more complex. The same procedure was adopted with the same parameters as used for the Tm1570 and TrmD domains separately. The unfolding trajectory of TrmD-Tm1570 starts from a flexible part of TrmD (its C-terminal region), with the first two rupture points corresponding to those seen in the TrmD domain unfolding, when the domain was unfolded alone. The knot cores of each domain remain in their native conformations, ***Figure 6***B(1-3). The end-to-end distances from the native state to point 1, and to point 2 are equal to 5.18 nm, 8.82 nm, and 25.85 nm. Equilibrium an explicit solvent, all-atom MD simulations indicated fluctuations in this value of up to 7.0 nm Next, the Tm1570 starts to unfold from its C-terminal side (rupture point 3, at 32.57 nm). Then, the unfolding of the TrmD domain from its N-terminal side takes place (rupture points 4 and 5), ***Figure 6***B(4-5). Between events number 5 and 6, the knot tightening process of the TrmD domain is observed. Surprisingly, there is no additional visible event between events 6 and 7, despite Tm1570 unfolding and its knot tightening between these steps, ***Figure 6***B(6-7).

***Figure 6***C displays the probability of native contacts during the stretching process. In agreement with the structural representations shown in panels A and B, these data reveal that the knot core in both domains are the final portion of the protein to unfold. The results highlight the role of the knotted region, which resists mechanical perturbation longer than other structural elements and show its importance in maintaining the local fold.

## Discussion

In this study, we employed various biophysical experimental techniques, MD simulations, and AI methods to investigate the thermal and thermodynamic stability, as well as the mechanical unfolding, of the double deeply knotted protein TrmD–Tm1570, along with its individual subunits, TrmD and Tm1570. The TrmD–Tm1570 protein serves as an ideal research model because each of its subunits also exists independently, forming the single deeply knotted proteins TrmD and Tm1570.

Our extensive MD simulations suggest that, to the best of our knowledge, coarse-grained models such as SMOG or UNRES – whether based on native contacts or not, and regardless of the type of potential used – are insufficient to fold the double-knotted TrmD–Tm1570 to the native conformation. While each domain can self-tie into its native knot, this process inhibits the knotting of the other domain. In contrast, native contacts are sufficient to fold the single-knotted TrmD and Tm1570 proteins, as has been shown previously for other knotted proteins from the SPOUT family using both computer simulations and experimental approaches.

Experimental studies contribute significantly to our understanding of the free energy landscape (***King et al., 2010***; ***Jackson et al., 2017***; ***Ziegler et al., 2016***; ***Zhang and Jackson, 2025***; ***Mepperi et al., 2025)***; however, some ambiguity in the interpretation of topology always remains. Setting that aside, the double-knotted TrmD–Tm1570 protein is challenging to study experimentally, as it rapidly aggregates in response to temperature changes and is produced only in small quantities. For this reason, we focused on mechanical pulling and thermal unfolding, using both coarsegrained simulations and optical tweezers experiments. Slow velocity stretching can provide valuable insights into the potential folding mechanism. Based on coarse-grained simulations, we identified four distinct unfolding and untying pathways for the TrmD–Tm1570 monomer. Notably, the most common pathway (Pathway #2) involves reversible unfolding and refolding of both domains without knot translocation. These pathways can be compared to mechanical unfolding and refolding scenarios in which the knots remain tied in the unfolded protein state.

Experimental unfolding and refolding of Tm1570, TrmD, and the TrmD–Tm1570 protein using single-molecule optical tweezers provided unique insights. Constant-velocity force–extension measurements for TrmD–Tm1570 and Tm1570 revealed multiple unfolding pathways; however, the most frequently observed pathway featured four and three major unfolding events, respectively, for the two proteins. For TrmD, unfolding and refolding traces showed a consistent unfolding pathway, with one smaller and one larger unfolding event with an increase in contour length. The measured changes in contour length for all three proteins suggest the presence of a structure or structures which are not unfolded under the applied forces at these constant stretching velocities, which we attribute to the presence of knots in the structure. Our measured changes in contour length are in good agreement with results from a structure-based coarse-grained model supported by an explicit-solvent all-atom model around the native state, enabling a detailed interpretation of the unfolding landscape and knot-tying mechanisms for double-knotted protein.

As part of this study, we will perform the folding of TrmD–Tm1570 in vitro in the presence and absence of molecular chaperones. While our current study demonstrates the unfolding and refolding processes, chaperones may play a critical role in vivo by preventing misfolding, reducing kinetic traps, or facilitating productive threading events

Moreover, consistent with previous degradation studies reported in ***da Silva et al. (2023)***, our data showed that TrmD is less mechanically stable than Tm1570 (***Figure 5*** and ***Table 1)***. This difference likely stems from the positioning of the knot core within each domain, which influences how the protein responds to mechanical forces. By integrating mechanical and thermal unfolding data, we characterized the stability landscape of the individual domains within the TrmD-Tm1570 fusion protein.

In summary, our study establishes that TrmD-Tm1570 represents a complex model system for investigating the stability and unfolding behavior of double-knotted proteins. We demonstrated that the two domains contribute differently to the overall stability of the double-knotted protein. We show the limitations of current coarse-grained models in the ability to simulate the folding of the double-knot fold motif in proteins, even though individual knots—regardless of their depth (***Piejko et al., 2020)*** or complexity—can (e.g. knot with 6 crossings, such as 6_1_ or 6_3_ (***Bölinger et al., 2010***; ***Sikora et al., 2023)***) self-tie within these models. Leaving that aside, it has been shown that chaperones can play an important role in knotting for one class of knotted proteins (***Mallam and Jackson, 2012)***; however, not all knotted proteins need assistance for rapid folding; some of them can self-tie quickly (***Wang et al., 2015)***. Thus, double-knotted proteins may represent a combination of these mechanisms, with one domain capable of self-knotting while the other depends on chaperone assistance.

In total, more than 1,200 proteins are predicted to contain double knots, including both globular and transmembrane (CaCA family) proteins. This work is therefore only the beginning of understanding how such motifs can arise.

## Methods and Materials

### Protein purification and expression

Three protein samples were prepared: TrmD, Tm1570 and the TrmD-Tm1570 protein, along with single-point mutations required to accommodate two cysteine residues, one at each terminus. For TrmD, these were introduced at positions M1C and T240C, for Tm1570 at positions G2C and L193C; and for the fusion at positions M1C and L433C. The appropriate mutants were prepared by PCR reaction through site-directed mutagenesis. Each protein was subcloned into *E. coli* BL21 DE3 RiL and grown in LB medium and induced using 1 mM IPTG. Proteins were purified by ion affinity chromatography and size exclusion chromatography. Further details are given in the ***da Silva et al. (2023)***.

### Differential scanning fluorimetry

DSF samples were prepared at increasing protein concentrations from 1 to 11 *μ*M, using a blank as a benchmark and Sypro Orange as the signal dye (Sigma-Aldrich). Measurements were performed on the CFX96 Real-Time PCR machine (Bio-Rad). The experiment began with 2 minutes of temperature equilibration at 22 ^°^C, followed by a temperature increase to 95 ^°^C at a rate of 0.5 ^°^C per 30 seconds, and fluorescence intensity (FRET channel) was measured at each step. All experiments were repeated at least 3 times to ensure reliability. Data analysis was conducted using custom scripts.

### Mechanical unfolding/folding with optical tweezer

Experimental mechanical unfolding was conducted by optical tweezers (C-trap, Lumicks, NL) as described previously(***Mondol et al., 2023)***, using 1.24 *μ*m antidigoxigenin and 1.21 *μ*m streptavidin-coated silica beads (Spherotech, USA). Measurements were performed at an average trap stiffness of 0.3 pN/nm and repeated unfolding/refolding cycles were collected at a constant approach and retraction velocities of 20 and 500 nm/s. Measurements were performed in 40 mM HEPES pH 8.0, with 500 mM NaCl and 10 % glycerol with the addition of an oxygen scavenger system comprising 1700 U/mL glucose catalase, 27 U/mL glucose oxidase, and 0.66 % glucose to minimize photodamage caused by oxygen free radicals. In order to prepare the proteins for optical tweezers measurements, samples of purified cysteine mutants of TrmD, Tm1750 and the double knotted TrmD-Tm1750 were functionalised with short single-stranded DNA oligonucleotides (for sequences refer to cite: Mondol et al., 2023) via a cysteine-maleimide reaction. This was done by first performing a buffer exchange step to remove the reducing agent that the sample was stored in (the storage buffer was the same as the measurement buffer with the addition of 1 mM TCEP). Then, each sample was incubated for 1h at room temperature with the DNA oligonucleotides at a 1.2:1 ratio of oligonucleotides per cysteine in the same buffer as the measurement buffer. Unreacted oligonucleotides were then removed via a size exclusion chromatography (SEC) step. Samples corresponding to monomeric TrmD, Tm1750 and the fusion protein with two oligonucleotides attached were aliquotted, flash frozen and stored for subsequent measurements. On the day of each measurement, samples were incubated for 1h at room temperature with double-stranded DNA handles (185 nm each) with a single-stranded DNA overhang complementary to the single-stranded DNA oligonucleotides on one end and either biotin or digoxigenin functional sites at the other end. Immediately prior to measurement, samples were incubated with anti-digoxigenin-coated beads for 15 minutes at room temperature. Finally, in the optical tweezers setup, tethers were formed by catching one anti-digoxigenin-coated bead and one streptavidin-coated bead and bringing them together.

The resulting force-extension traces were analysed using the worm-like chain (WLC) model to fit the protein unfolding events and the extensible worm-like chain (eWLC) model to fit the DNA stretching, as described before(***Mondol et al., 2023)***, with an DNA contour length of 370 nm, a DNA persistence length of 20 nm, a temperature of 293 K, and the elastic stretch modulus of the DNA (K) set to 500 pN. The measurements were performed with a sampling rate of 78 kHz, giving a time resolution of 13 ms.

### Mechanical unfolding simulations

The mechanical unfolding process was investigated by MD coarse-grained (CG) simulations with the structure-based C*α* model (SBM-C*α*) (***Clementi et al., 2000)***. The SBM-C*α* defines the potential in such a way that the minimum energy value is attributed to the protein’s native structure (***Cieplak and Sułkowska, 2011)***. All simulations were performed using the Gromacs 4.5.5 software package(***Van Der Spoel et al., 2005)*** with a leapfrog integrator. Input files were generated with the SMOG server(***de Oliveira Jr et al., 2022)***. To maintain a constant temperature, the Berendsen thermostat algorithm(***Berendsen et al., 1984)*** was employed with a coupling time constant equal to 1 ps. Protein simulations were initiated in their native configurations with 5 x 10^8^ steps, and the timestep is equal to 0.5 fs. Configurations were saved every 1000 steps. The fraction of native contacts (Q) is the reaction coordinate used to monitor the simulations. Two residues, *i* and *j*, are interacting when the distance between their C*α* atoms is shorter than 1.2d_ij_. d_ij_ represents the distance between C*α* residues in the native structure, where *j > i* + 3. The contact map was defined using the Shadow Contact Map software(***Noel et al., 2012)***.

In the stretching simulations conducted under constant speed, *ν*, the C-terminus of the protein is attached to harmonic elastic springs and it moves at a speed, *ν* = 0.05 Å/*τ*. In the course of simulations, the N-terminus of the protein was fixed and the C-terminus was subjected to stretching forces. The TrmD-Tm1570, TrmD, and Tm1570 present the same mutations as reported in the *Experimental section* and the residues to which stretching forces were applied are located in the C-terminal region: C433, C240, and C193 for the TrmD-Tm1570, TrmD, and Tm1570 proteins, respectively.

Time in the coarse-grained simulations is expressed in units of the Lennard-Jones time scale 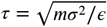, which for our model corresponds to 3.2 ps. Pulling speeds of 0.05, 0.10, and 0.15 Å/*τ* therefore correspond to 1.6–4.7 m/s in real units. The simulations are performed in an implicit-solvent regime and do not include explicit hydrodynamic friction, and therefore the absolute timescales do not correspond directly to experiment. Instead, we ensured that the unfolding pathways and force–extension profiles were consistent across simulated pulling speeds, allowing a qualitative comparison with the experimental unfolding mechanisms.

### Thermal unfolding simulations

#### Coarse-grained model

The thermal unfolding MD simulations were conducted to investigate the unfolding pathway of the protein upon increasing temperature. The MD simulations were carried out in a range of temperatures, *T*, from 0.9 to 1.6 (reduced units), with Δ*T* = 0.04, for a total of 17 different temperatures. Additionally, 200 simulations were performed at the temperature where a slow transition from the native to the unfolded state was observed. Thermal unfolding was investigated by monitoring the fraction of native contacts (Q). All the parameters are the same as those used in the *Mechanical Unfolding Simulations*.

#### All-atom model

The proteins were protonated in pH 7.0 using the PDB2PQR server and neutralized by adding sodium cations and chloride anions. The explicit water TIP3P model was used. The MD simulations were performed in the GROMACS 2023.3 software package with the CHARMM36 force field. All performed production runs shared common first steps of the system preparations, including energy minimization and relaxation. Equilibration prior to the production runs included an NVT ensemble and two phases of the NPT ensemble. For the temperature equilibration, a V–rescale temperature coupling was used. The temperature, 296 K, was selected to provide the best possible accordance with the existing experimental data.

### Architecture of the Self-Organization Map - SOM

A SOM is a valuable tool for exploring, understanding, and classifying data points. A SOM or Kohonen neural network (NN) is an unsupervised NN(***Kohonen, 1982, 1998)***, it can learn from unlabeled data by processing it and transforming it into a 2D map. The map preserves the relationships between the original data points, meaning similar data points are mapped close together(***Kohonen, 2013)***. This technique is useful for visual inspection of complex datasets and identification of patterns or clusters that might be difficult to detect in the high-dimensional space. SOM is based on three key principles: 1-Initialization; 2 - Competition and Cooperation; and 3 - Iterative Learning.

1. <u>Initialization</u> - (I) Dimensions of the SOM: the size of the SOM grid is equal to 4, a 4×4 grid is equal to 16 nodes; (II) Input Length: the number of features in the input data; (III) Neighborhood Radius: a function that weights the neighborhood of a position in the map, the neighborhood function is set to be Gaussian. (IV) Learning rate: this defines how much the SOM weights are updated during each iteration of training, the learning rate is set to be equal to 0.5. (V) Randomly initialize the weights of each neuron.
2. <u>Competition</u> - Given the input data, the SOM performs a competition among all units. Each unit calculates the distance between its weight vector, *w*_*i*_. The input vector, *x*, using a Euclidean distance, *argmin*_*i*_ ‖*x* − *w*_*i*_ ‖. The unit with the closest weight vector (th winner node) is declared the Best Matching Unit (BMU).
3. <u>Iterative Learning</u> - Once the BMU is identified, its weight vector and the weights of its neighboring units are adjusted to become more similar to the input data. This update is called adaptation and involves a learning rate, *η*, that determines the magnitude of the update as follow in Equation 1:

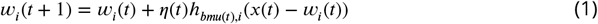

where *w*_*i*_(*t*) is the weight vector of neuron *i* at time step *t. η*(*t*) is the learning rate at time step *t. h*_*bmu*(*t*) *i*_ is neighborhood function value for neuron *i* relative to the BMU at time step *t. x*(*t*) is the input vector at time step *t*.

Unlike other neural networks that count on error correction learning, SOM utilizes a competitive learning strategy. Giving the input data, SOM neurons compete via a “Winner-Takes-All” (WTA) mechanism. The winning neuron, along with its neighbors in the SOM grid, adjusts its weights to better represent the input. Through this competitive mechanism, neighboring neurons display similar patterns, while neurons further away inhibit distant units. This interplay between competition and cooperation allows the SOM to self-organize and effectively capture the similarity of the data, see the SOM illustration in ***Figure 7***.

**Figure 7.**
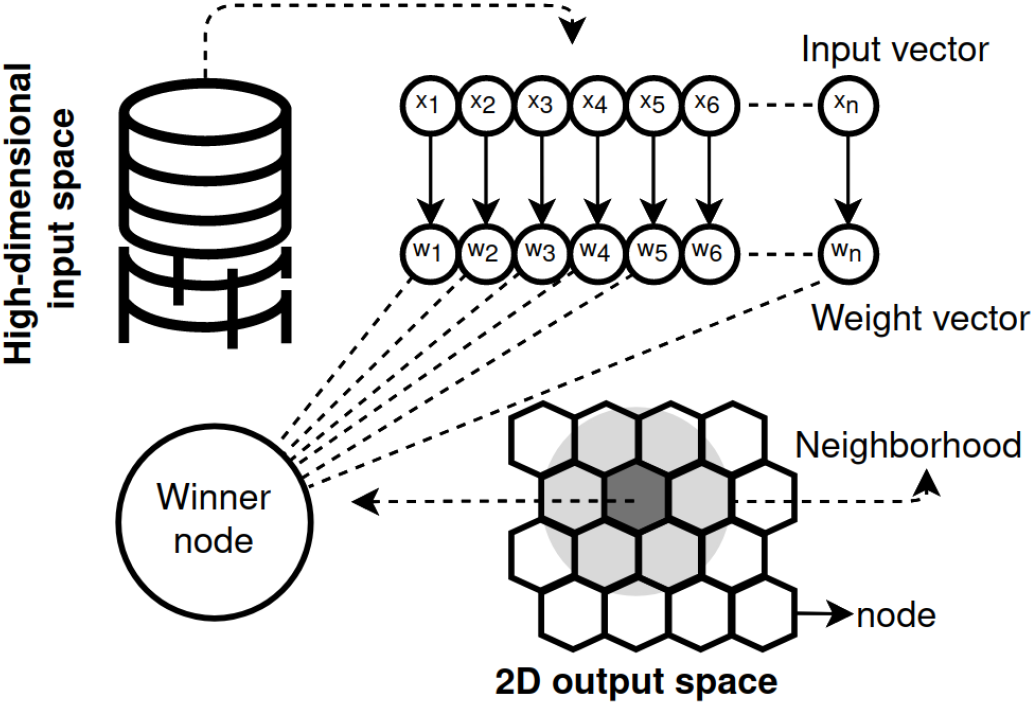
Self-Organizing Map (SOM) representation. The high-dimensional input space represents the input dataset, fraction of native contacts (*Q*). Each input vector, *x*_*i*_, is associate with weight vector, *w*_*i*_. The neurons are in a *k* × *ℓ* lattice, SOM grid, 2D output space. The SOM utilizes a geometric layout, typically rectangular or hexagonal, where each individual unit within the grid is a neuron.

## Acknowledgments

This work was supported by the National Science Centre (#UMO-2018/31/B/NZ1/04016 to JIS) and Excellence initiative – Research University (BOB-661-1737/2025 to FBS). We would like to thank Maciej Lukasiewcz for help with DSF approach. The author(s) would like to acknowledge the contribution of the COST Action CA21169, DYNALIFE, supported by COST (European Cooperation in Science and Technology). Declarations of interest: none.

## Author Contributions

F.B.S.: conceptualization, data curation, formal analysis, investigation, methodology, visualization, writing—original draft, funding acquisition, writing—review & editing. S.N.: conceptualization, data curation, formal analysis, investigation, methodology, visualization, writing—review & editing. I.L.: data curation, formal analysis, investigation, methodology, M.F. and M.S.: data curation, formal analysis, investigation. L.M.S.: data curation, formal analysis, investigation. K.T.: conceptualization, data curation, formal analysis, investigation, methodology, visualization, writing—review & editing. J.I.S.: conceptualization, formal analysis, investigation, methodology, visualization, funding acquisition, project administration, supervision, writing—review & editing.

## Appendix 1

### Differential scanning fluorimetry measurements

Samples were prepared in the increasing concentrations of protein from 1 *μ*M to 11 *μ*M with a blank as a benchmark and Sypro Orange (5x) as the signal die. The experiment started with 15 minutes of temperature equilibration at 25 ^°^C after which the temperature was raised to 90 ^°^C at a rate of 1 ^°^C per minute. Data analysis was conducted using MicroCal DSC-ORIGIN analysis software and self-written scripts.

### Double knotted TrmD-Tm1570 is as stable as single-knotted protein TrmD – thermal denaturation approach

To investigate the thermal stability of TrmD, Tm1570, and TrmD-Tm1570, we employed DSF (Differential Scanning Fluorimetry measurements), a sensitive technique that measures changes in protein fluorescence upon thermal denaturation. The DSF analyses were conducted at different protein concentrations, from 1 to 10 *μ*M, Figure S1; for each melting curve, we determined the melting temperature (*T*_*m*_) of the proteins, ***Figure 1***. Based on data from other publications about SPOUT, we assume that we observe denaturation of monomers, even though TrmD, Tm1570, and TrmD-Tm1570 exist as dimers. The *T*_*m*_ of TrmD and Tm1570 increases respectively from 48.5 to 64.0^°^C; 59.5 to 73.5^°^C as protein concentration increases. *T*_*m*_ for fusion is below *T*_*m*_ for Tm1570. The melting temperature we measured for TmrD correlates with the expected range for proteins belonging to the SPOUT family. This suggests that Tm1570 exhibits significantly higher thermal stability compared to TrmD; however, the addition of TrmD to Tm1570 – a fusion protein – destabilizes the protein. That is, a doubly knotted protein is possibly not thermally more stable than a single one; local properties of Tm1570 protein have a greater impact than the topology itself. Consequently, Tm1570 emerges as the most stable protein within this family.

The estimated melting temperature of TrmD-Tm1570 remains relatively stable, from 58.5 to 62 ^°^C, across the concentration range, suggesting that concentration does not significantly affect its thermal stability (***Figure 1*** - top panel) or we observed aggregation. The single knotted proteins (Tm1570 and TrmD) exhibit concentration-dependent increases in *T*_*m*_, indicating that higher concentrations promote their thermal stability. However, no additive effect is observed in TrmD-Tm1570, despite Tm1570 being thermodynamically more stable than TrmD. TrmD and TrmD-Tm1570 display nearly identical *T*_*ms*_ at concentrations between 5 and 10 *μ*M. However, these results should be treated with caution, as the protein has a very high tendency to aggregate.

By setting up the temperature at 60 ^°^C, we calculated the unfolded fraction for each protein in different concentrations, ***Figure 1*** – bottom panel and ***figure Supplement 1***, lower panel. We observe that the increase in *T*_*m*_ and decrease in unfolded fraction for TrmD and Tm1570 at lower concentrations suggest that these proteins are stabilized by intermolecular interactions at higher concentrations. Additionally, the Tm1570 appears to confer some degree of concentration-dependent stabilization to TrmD-Tm1570, as evidenced by the monotonic decrease unfolded fraction of TmD-Tm1570 compared to TrmD.

**Appendix 1—figure 1.**
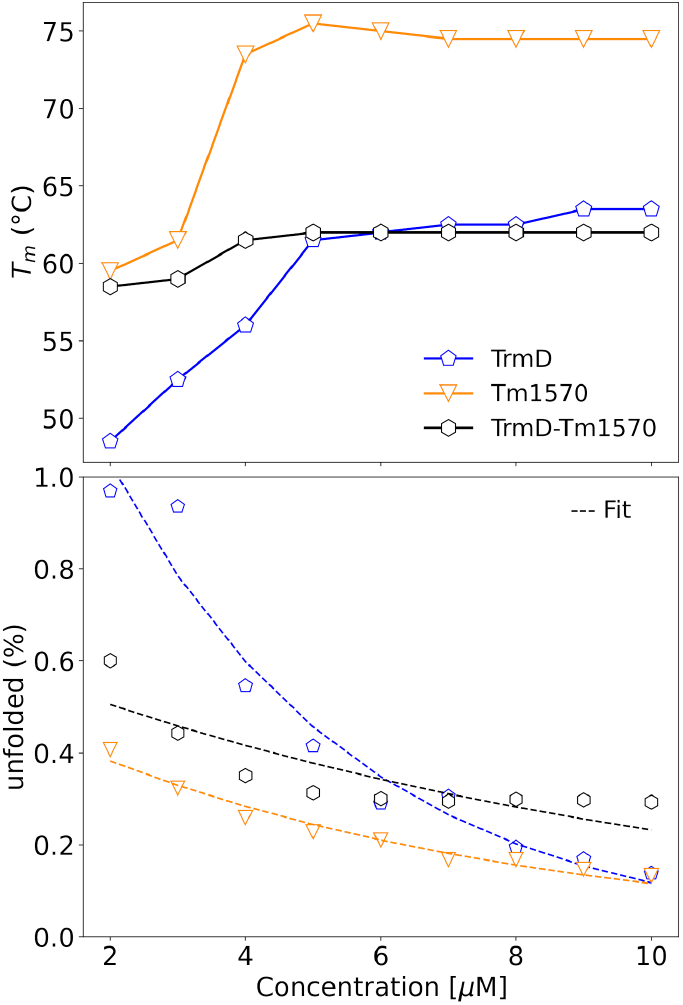
Melting temperature (*T*_*m*_) under different protein concentrations (1-10 *μ*M). TrmD, Tm1570, and TrmD-Tm1570 are shown in pentagon, triangle down, and hexagon markers in colors blue, orange, and black, respectively. The fluorescence melting curves measure by DSF in Figure S1. **Appendix 1—figure 1—figure supplement 1**. Thermal Denaturation of TrmD, Tm1570, and TrmD-Tm1570.

## Appendix 2

### Description of rare events observed

This section presents nine rare events observed in our simulations, Figures S1A-I, associate with the main ***Figure 3***, divided into three groups: 1) complete unfolding process, Figures S1(A,B, and H); 2) partially unfolding event, Figures S1(C, F, and I); and 3) cyclic process, Figures S1(D, E, and G).

1. **Complete unfolding process:** In these group of simulations, Figures S1(A, B, and H), the refolding process of TrmD-Tm1570 is observed. As mentioned in the main manuscript, the refolding process occurs once the knot remains at the same native position. In Figure S1A, both domains unfold, 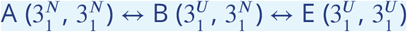, following by the Tm1570 untied process, 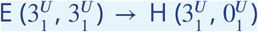, and refold, unfold, and untied process to TrmD domain, 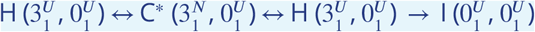. In Figure S1B. the unfolding process is similar with path 1 reported in the main manuscript. However, it starts with unfolding-refolding event in Tm1570 domain, 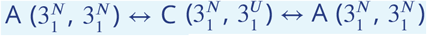. In Figure S1H, the complete unfolding process is similar to the path 4 presented in the main manuscript. Despite that after Tm1570 untied event, the TrmD domain refold to its native state, unfold and untied, 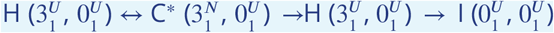.
2. **Partially unfolding process:** In this group of simulations we cannot observe the complete unfolding process due to the simulations time giving as an input. However, it may be possible to observe with the simulation time extension. In all cases the refolding process is observed, but only in Figure S1C, the refolding event occurs without previously untied process. The process in Figure S1C is represented by a cyclic process initially and then a TrmD untied event, 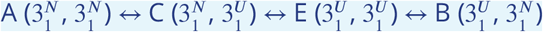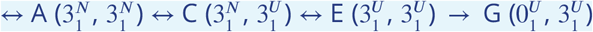. In Figures S1F and S1I, the refold event of TrmD and Tm1570 are observed after untied process of Tm1570 and TrmD, respectively.
3. **Cyclic process:** These events are represented by the unfolding and refolding process without untied or complete unfolding events. In Figure S1D, the unfolding process starts by the TrmD domain, following by its refold and finally by the Tm1570 unfolding, 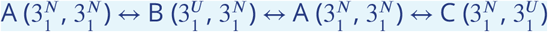. On the other hand, in Figure S1G, the unfolding process starts by the Tm1570 domain, following by its refold and finally by the TrmD unfolding,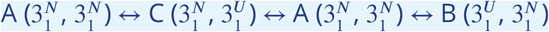.

**Figure 3—figure supplement 1.**
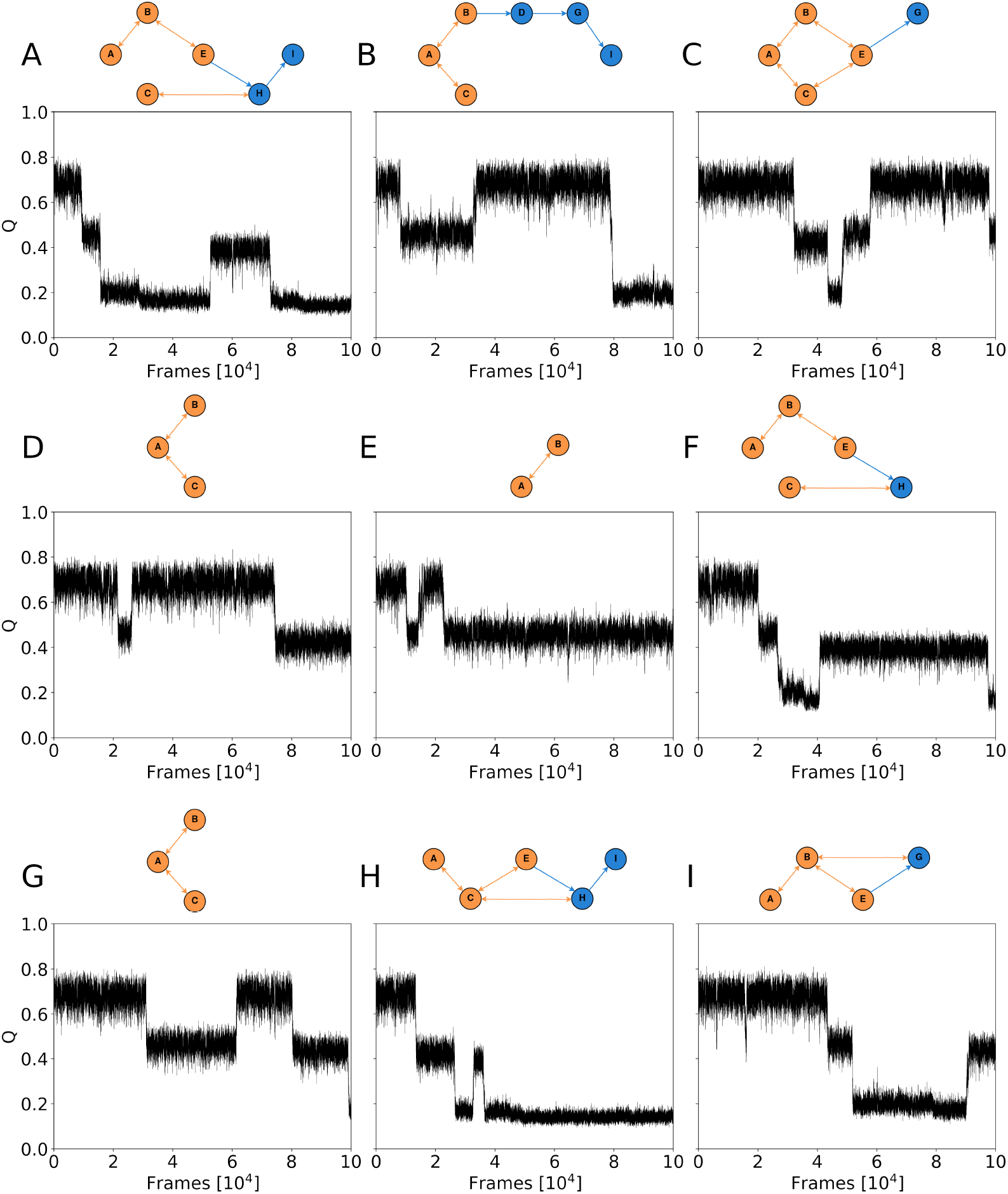
Unfolding trajectories observed less frequently, rare events. **Panel A –**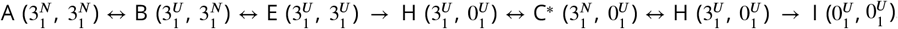. **Panel B –** 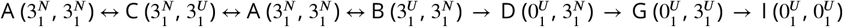. **Panel C –** 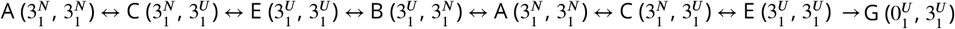. **Panel D –** 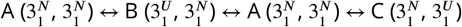. **Panel E –** 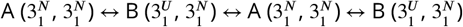. **Panel F –** 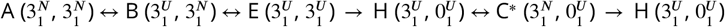. **Panel G –** 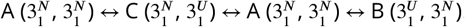. **Panel H –** 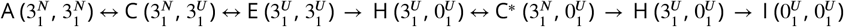. **Panel I –** 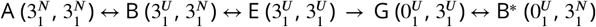.

**Figure 3—figure supplement 2.**
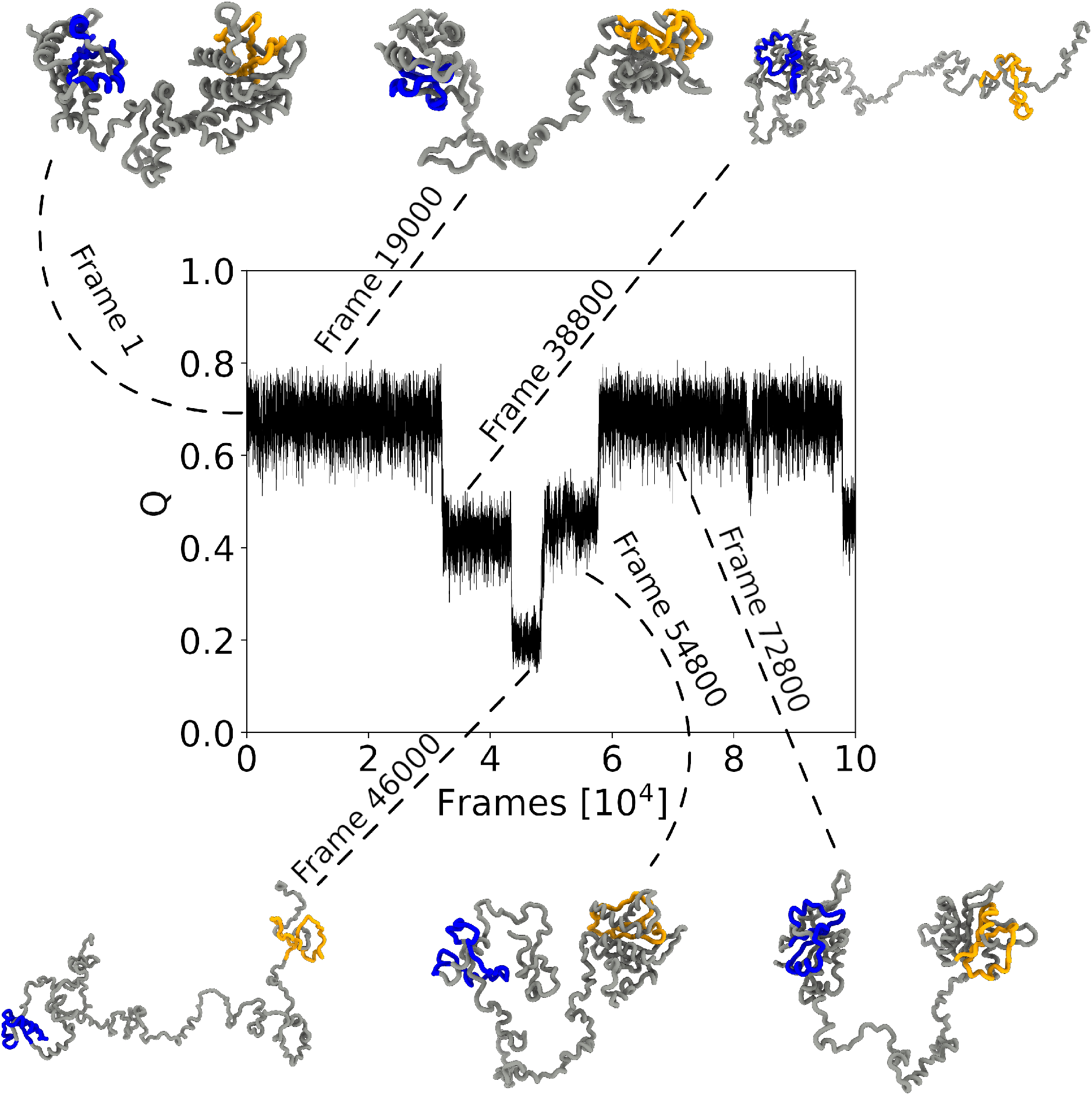
Refolding dynamics of a fusion protein, visualized through snapshots at key time steps (frames 1, 19000, 38800, 46000, 54800, and 72800). The knot cores, localized within residues 85-129 (TrmD domain, blue) and 352-397 (Tm1570 domain, yellow), remain invariant during the refolding process.

**Figure 3—figure supplement 3.**
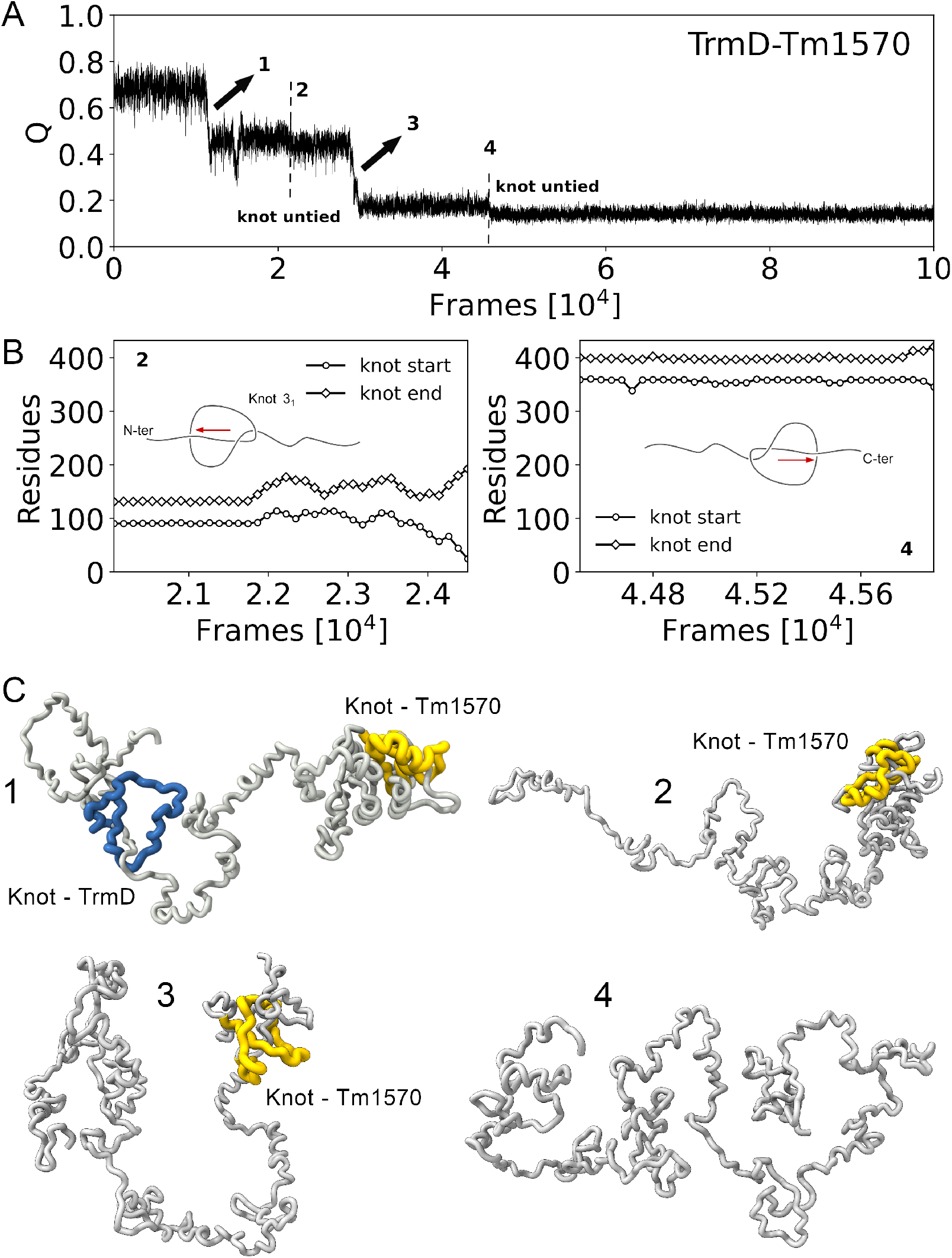
(A) Fraction of native contacts (Q) during constant-velocity pulling simulations. Four unfolding events are annotated (1–4), along with two distinct knot untying transitions (2 and 4). (B) Knot position for events 2 and 4. The left plot shows the movement of knot from TrmD domain during event 2, around frame 2.2 × 10^4^. The right plot shows the corresponding knot dynamics from Tm1570 domain during event 4, around frame 4.56 × 10^4^. (C) Snapshots from the simulation of the fusion protein at four key time points corresponding to events 1–4. Knotted regions are highlighted in blue (TrmD) and yellow (Tm1570), illustrating the untying events of both domains and the transition from a double-knotted to a fully unknotted state.

**Figure 4—figure supplement 1.**
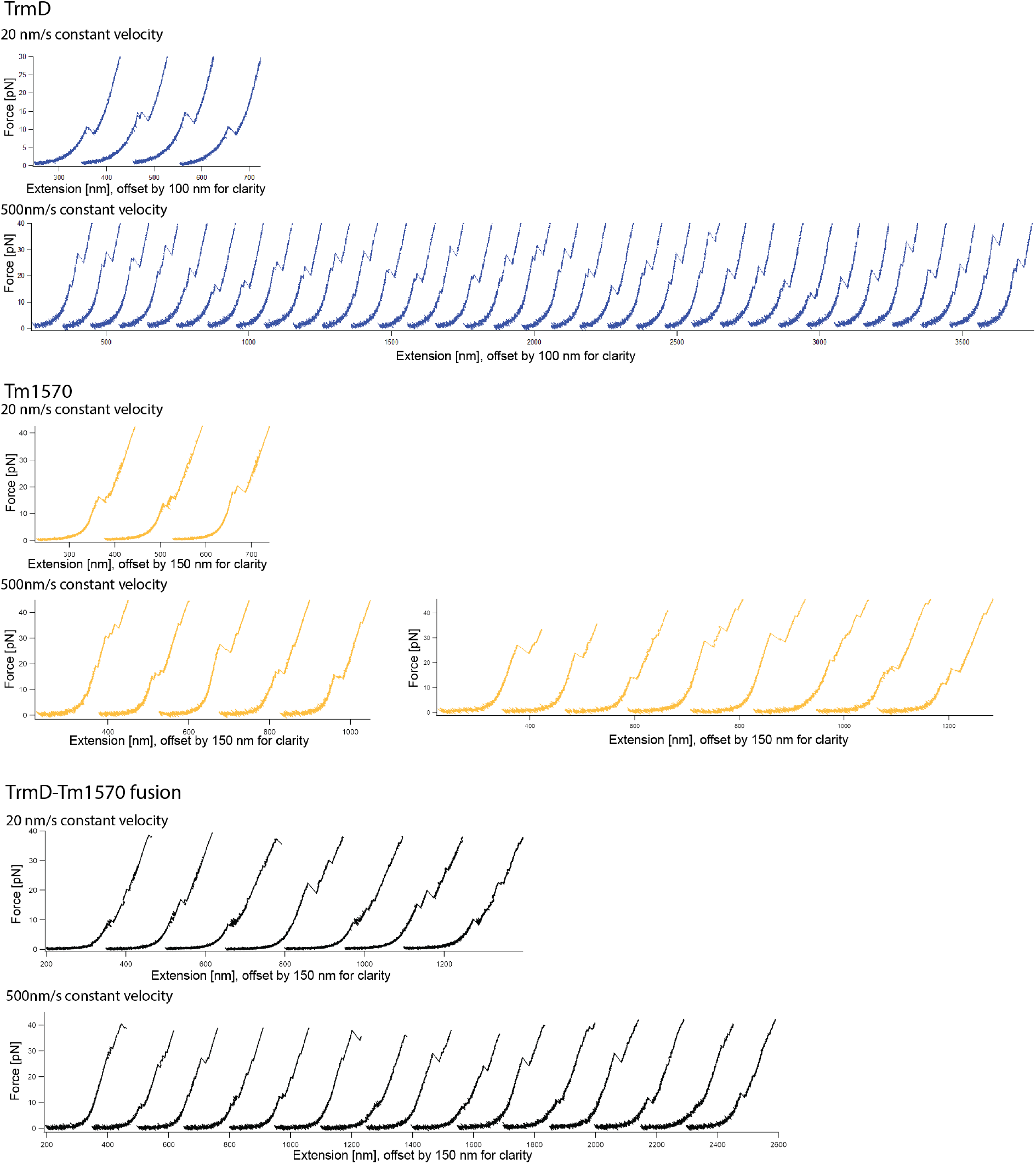
Experimental unfolding of Tm1570, TrmD and the Tm1570-TrmD Fusion using single-molecule optical tweezers. In each trace the unfolding is colored (dark blue for TrmD, orange for Tm1570, and black for the TrmD-Tm1570). The traces were measured at a constant velocity of 20 and 50 *nms*^−1^.

**Figure 5—figure supplement 1.**
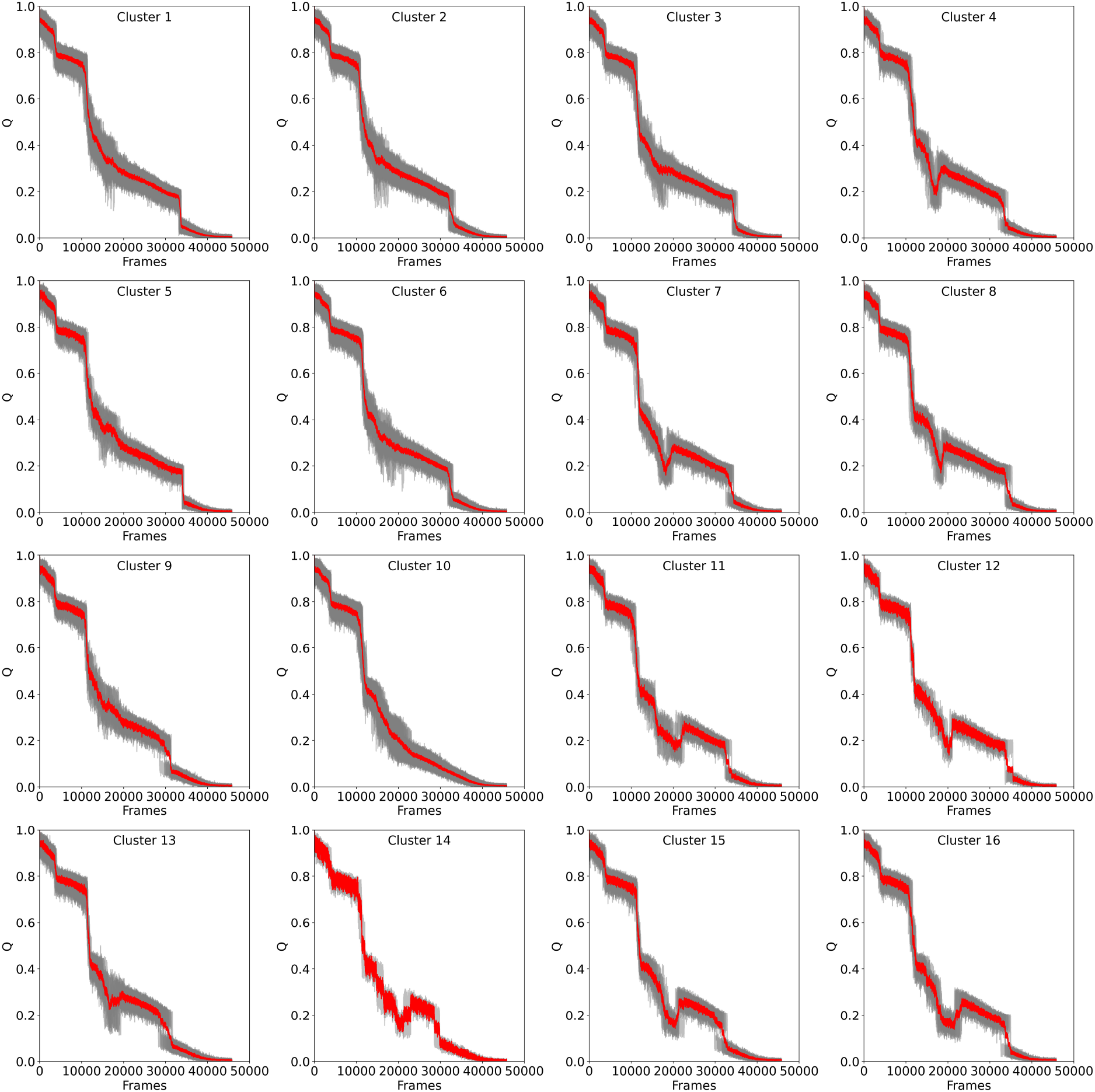
Clustering of the unfolding trajectories for Tm1570 obtained via SOM analysis. Each subplot (Clusters 1–16) displays the average unfolding profile (red line) of a cluster, plotted as the fraction of native contacts (Q) versus simulation frames. Grey traces represent individual trajectories within each cluster.

**Figure 5—figure supplement 2.**
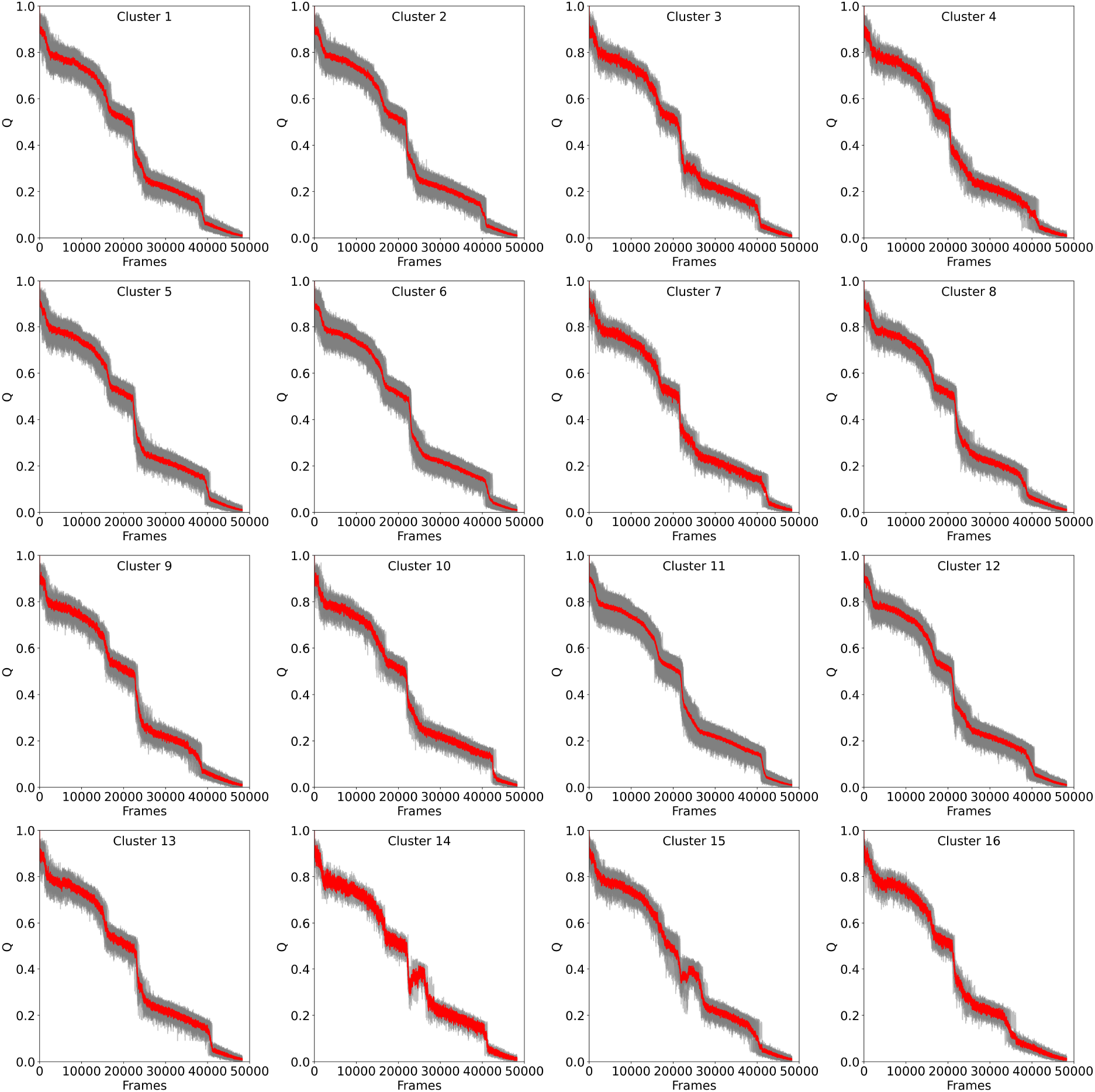
Clustering of unfolding trajectories for TrmD obtained via SOM analysis. Each subplot (Clusters 1–16) displays the average unfolding profile (red line) of a cluster, plotted as the fraction of native contacts (Q) versus simulation frames. Grey traces represent individual trajectories within each cluster.

**Figure 5—figure supplement 3.**
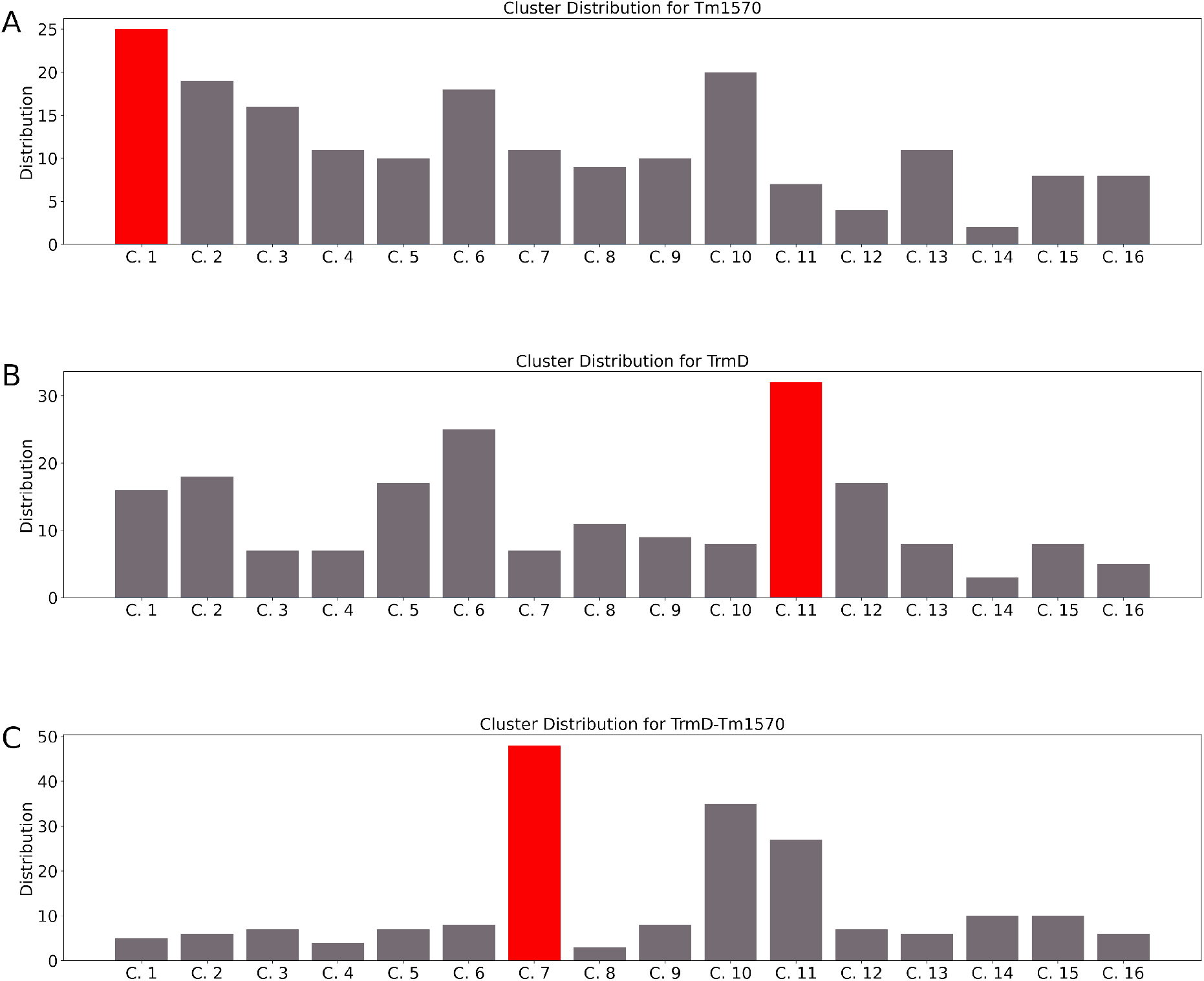
Distribution of simulations for each protein across clusters. Panels A, B, and C show Tm1570, TrmD, and TrmD-Tm1570, respectively. The most populated cluster for each protein is highlighted in red: cluster 1 for Tm1570, cluster 11 for TrmD, and cluster 7 for TrmD-Tm1570.

**Figure 5—figure supplement 4.**
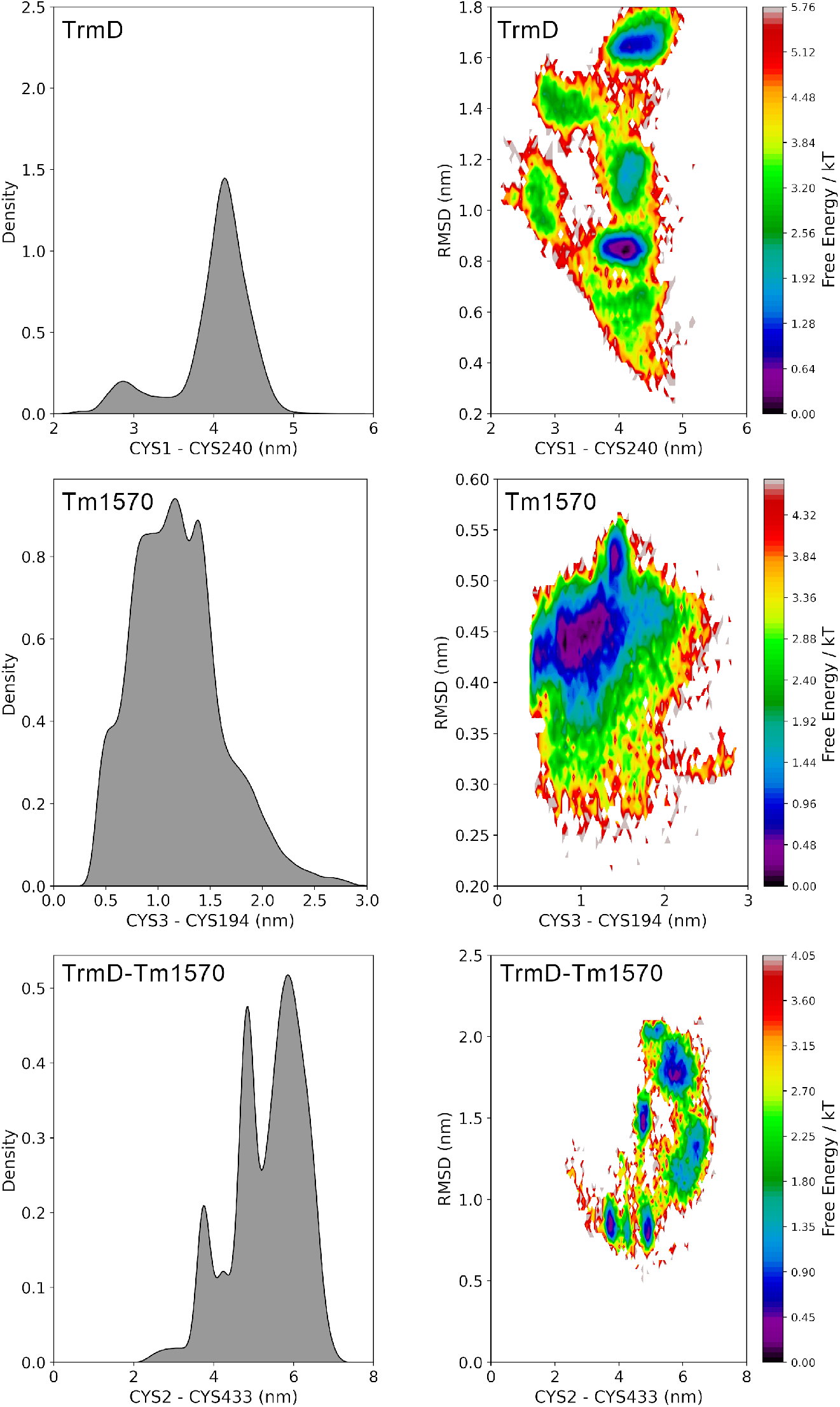
Probability densities and free energy landscapes for TrmD, Tm1570, and the TrmD–Tm1570 proteins. Left panels: probability density of end-to-end distances between cysteine residues. Right panels: free energy surfaces as a function of end-to-end distances and RMSDs, with color indicating free energy in units of kT. Probability density and free energy were computed using the package PyEMMA in python version 3.8.8. Details about the MD simulation in *Methods and Materials*.

**Figure 6—figure supplement 1.**
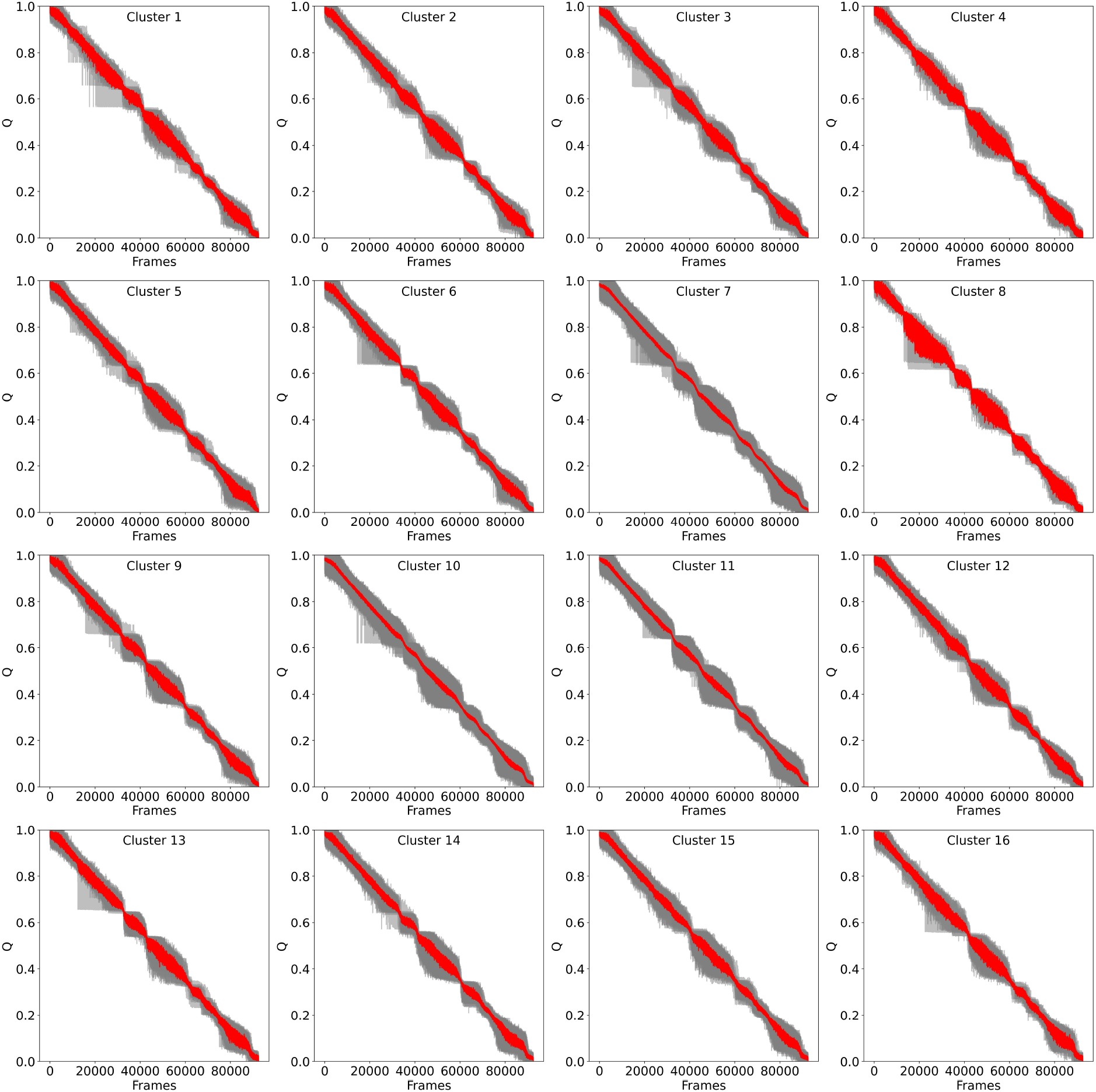
Clustering of unfolding trajectories for Tm1570-TrmD obtained via SOM analysis. Each subplot (Clusters 1–16) displays the average unfolding profile (red line) of a cluster, plotted as the fraction of native contacts (Q) versus simulation frames. Grey traces represent individual trajectories within each cluster.

**Appendix 1—figure 1—figure supplement 1.**
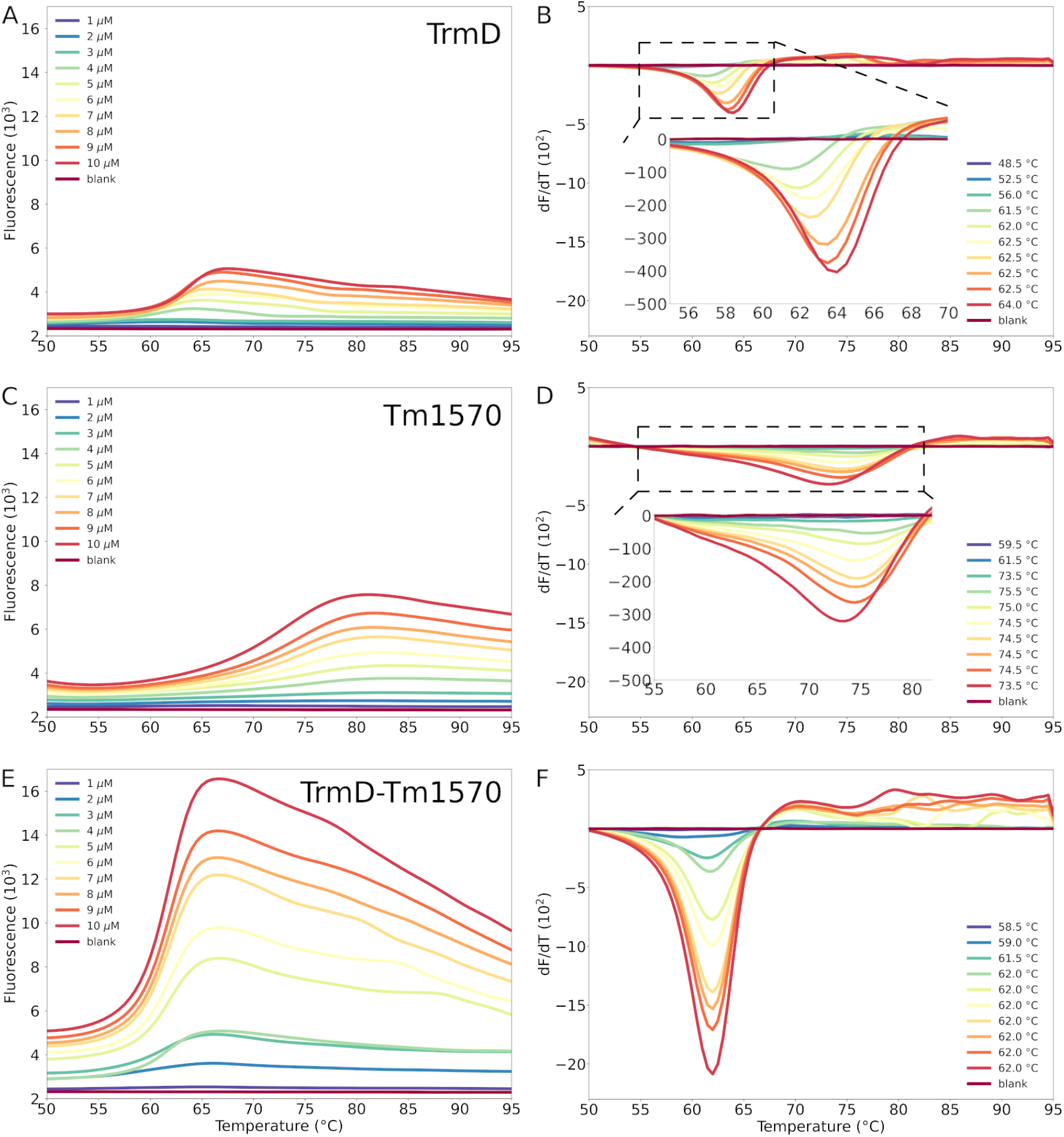
Thermal Denaturation of TrmD, Tm1570, and TrmD-Tm1570. A, C, and E - Fluorescence melting curves of TrmD, Tm1570, and TrmD-Tm1570, obtained by monitoring the decrease in fluorescence intensity as the temperature increases. The analysis were conducted under different protein concentrations, from 0 (blank) to 10 *μ*M. The decrease in fluorescence intensity is due to the protein unfolding. B, D, and F - First derivative of the melting curves, highlighting the inflection point corresponding to the melting temperature (Tm). Results shown here are from one replicate.

